# Determining sex differences in drug combinations targeting aortic valve myofibroblast activation using an artificial intelligence derived platform

**DOI:** 10.1101/2024.10.02.615251

**Authors:** Brandon J. Vogt, Peter Wang, Megan Chavez, Peng Guo, Edward Kai-Hua Chow, Dean Ho, Brian A. Aguado

## Abstract

Aortic valve stenosis (AVS) is a sexually dimorphic disease where aortic valve leaflets develop fibrosis and calcification, leading to heart failure if untreated. Sex differences in AVS progression depend on valvular interstitial cells (VICs) activating to myofibroblasts that drive aberrant extracellular matrix remodeling. To date, no treatment strategies have leveraged cellular sex differences to determine drug combinations that effectively target VIC myofibroblast activation. Here, we harnessed IDentif.AI, an artificial intelligence (AI)-derived drug optimization platform, to optimize sex-specific synergistic drug combinations that may prevent and reverse VIC myofibroblast activation on hydrogel biomaterials. The results reveal that anti-fibrotic drug efficacy and combinatorial interactions are dependent on cell sex. This study provides a framework for developing clinically relevant AVS treatment strategies through the integration of high-throughput hydrogel cell culture platforms and AI-driven drug optimization. The workflow towards designing targeted AVS drug combinations may help accelerate AVS drug development for male and female patients and address health disparities in AVS treatment outcomes.

## Introduction

Aortic valve stenosis (AVS) impacts nearly 1 in 8 adults over the age of 75 and is characterized by progressive fibrosis and calcification of the aortic valve leaflet^1,2,3^. Valvular fibro-calcification impedes hemodynamic function of the left ventricle, culminating in heart failure and an expected 2–5-year mortality rate if untreated^4^. The current standard of care for AVS is surgical or transcatheter aortic valve replacement^5,6^. However, not all patients are eligible for these procedures and patients undergoing valve replacement procedures may experience restenosis of the implanted aortic valve^7,8^. To address the limitations of valve replacements, effective drug treatments may serve as an alternative to surgical intervention.

Drug treatments for AVS have resulted in little to no efficacy in large-scale clinical trials^9,10,11,12^. The failure of prior drug treatments for AVS is multifactorial, but is likely due in part to the historical failure to account for the known sex differences in AVS pathophysiology^13,14^. Indeed, sex as a biological variable has emerged as a predictor of AVS incidence, valve phenotype, response to treatment, and clinical outcomes^15,16,17,18^. For example, moderately-diseased female AVS patients broadly experience higher rates of valvular fibrosis relative to disease-matched male patients who instead experience increased calcification^17^. These sex differences lead to suboptimal treatment recommendations and outcomes, such as increased risk for female patients undergoing valve replacement procedures and undertreatment of moderately diseased female AVS patients^15,19,20,21^.

Sex differences are also observed on a cellular level, where female valvular interstitial cells (VICs), the native fibroblast of the aortic valve that comprises the majority of the aortic valve leaflet, are more prone to activating to a myofibroblast phenotype than male VICs, partially due to sex chromosome linked genes^22,23^. Activated myofibroblasts present in female valve tissue have been associated with increased extracellular matrix remodeling, collagen deposition, and fibrosis in female patients^17,24^. Sex differences in VIC phenotypes have been recapitulated *in vitro* using poly(ethylene glycol) (PEG) hydrogel biomaterials that mimic the stiffness of the valvular extracellular matrix^22,25,26^. Critically, culturing VICs on PEG hydrogels preserves the known female bias in myofibroblast activation, and thus has the potential to improve the clinical translatability of *in vitro* drug screening platforms that frequently rely on tissue-culture polystyrene (TCPS)^27^.

Another barrier to developing successful drug treatments for AVS includes accounting for the redundancies in myofibroblast signaling pathways that often render individual drug monotherapies ineffective at low doses. Therefore, there is an urgent need to properly combine multiple drugs and pinpoint optimal dose ratios for enhanced treatment outcomes^28,29^. However, optimizing effective drug combinations and their dose ratios is a challenging task as the parameter space consisting of all possible drug combinations scales exponentially with the number of drugs and doses tested in combination. To address this challenge, multiple approaches including machine learning and artificial intelligence (AI)-powered solutions have been explored^30,31,32,33,34,35,36^. In this study, IDentif.AI, an AI-derived drug optimization platform, was implemented to determine effective drug combinations that may slow or halt AVS progression. This platform interrogates the drug parameter space by correlating the input drug combinations and their corresponding biological response via an AI-discovered second order quadratic series and, subsequently, provides a ranked list of predicted biological responses for all possible combinations in the parameter space^37,38^. The AI-discovered relationship was originally determined via modeling of cellular responses to therapeutics using neural networks, which found that the relationship can be accurately modeled by a simple quadratic function^36^. Furthermore, IDentif.AI pinpoints optimal drug combinations independent of drug mechanisms of action and relies solely on prospectively generated data, instead of pre-existing information or big data. This small data-based approach has been validated in multiple disease models and led directly to clinical studies^39,40,41,42,43^. As such, IDentif.AI has emerged as a useful tool for determining drug combinations to modulate a wide array of cellular phenotypes and optimize therapeutic doses for multiple disease indications spanning from infectious diseases to oncology^44,45,46,47,48,49^.

In this study, we harnessed IDentif.AI to interrogate a pool of eight drug candidates that target myofibroblast signaling pathways and pinpoint effective drug combinations to inhibit myofibroblast activation distinctly in male or female VICs. We hypothesized that female VICs would be more resistant to individual anti-fibrotic drugs due to interactions between X-chromosome linked genes and myofibroblast signaling pathways^22^. Furthermore, we proposed that male and female VICs would require distinct drug combinations to effectively inhibit myofibroblast activation. To test our hypotheses, we developed a high-throughput hydrogel cell culture platform that is compatible with large-scale drug screening. We seeded male or female VICs onto this hydrogel platform, exposed them to combinations of eight inhibitors of interest at sex-optimized doses, and measured myofibroblast activation via alpha-smooth muscle actin (αSMA) immunofluorescent staining to develop predictive models of sex-specific myofibroblast inhibition. The IDentif.AI-designed drug combinations for male and female VICs were experimentally validated *in vitro* to confirm therapeutic efficacy. Sex-specific drug combinations were identified and validated by leveraging sex differences in drug synergies to demonstrate the efficacy of a precision medicine approach for AVS treatment. The integration of the high-throughput hydrogel cell culture platform and IDentif.AI may lay groundwork towards the development of AVS-specific therapeutics.

## Results

### Anti-fibrotic drug efficacy is dependent on cell sex

Based on established models of VIC myofibroblast signaling networks^29^ and prior work probing the effects of various inhibitors on myofibroblast activation, eleven anti-fibrotic drugs were selected for investigation (Supplementary Table 1). As an initial screen, we assessed the ability of each inhibitor to reduce myofibroblast activation in male and female VICs cultured on TCPS by quantifying the percent reduction in αSMA gradient mean intensity (GMI) through immunofluorescent staining (Supplementary Fig. 1). We found that eight out of the eleven anti-fibrotic drugs achieved meaningful reductions in αSMA GMI (>25%) in both male and female VICs in the dosing range of 10^−7^-10^3^ μM (Supplementary Fig. 2). Therefore, eight inhibitors were selected for drug combination optimization (Y-27632, LY294002, H1152, SB203580, Irosustat, TM5441, Losartan, and SD-208). The selected inhibitors were then tested at five doses on TCPS to fit drug response curves and determine an effective therapeutic dose for male and female VICs respectively (Figs. 1A-H). Six out of the eight inhibitors were more effective in male VICs relative to female VICs as quantified by a lower EC_20_ value (Fig. 1I). Y-27632 and Losartan were significantly more effective in male VICs relative to female VICs, and no inhibitors were significantly more effective in female VICs (Supplementary Fig. 3).

**Fig. 1.**
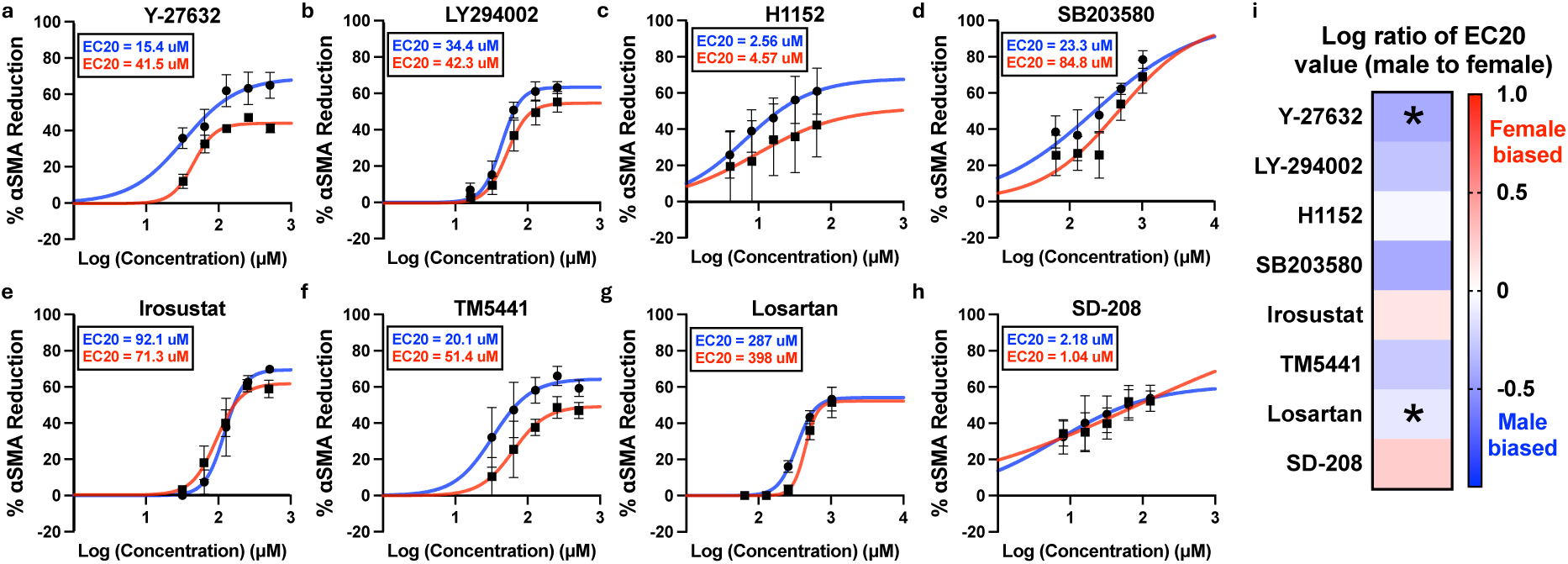
Drug response curves for anti-fibrotic drugs inhibiting male and female VIC myofibroblast activation on TCPS. **a-h,** Percent αSMA reduction in male VICs (blue) and female VICs (red) cultured on TCPS with indicated doses of **a** Y-27632, **b** LY294002, **c** H1152, **d** SB203580, **e** Irosustat, **f** TM5441, **g** Losartan, and **h** SD-208. **i,** Heat map showing the log ratio of male EC_20_ to female EC_20_ for each inhibitor. Data is plotted as mean ± standard error of the mean (N = 3 biological replicates). The best fit line shown was generated using a nonlinear regression curve fit with GraphPad Prism. Statistical significance was determined by an unpaired two-tailed t-test with Welch’s correction and indicated as *=P<0.05.

### Developing a hydrogel platform for high-throughput drug screening

Prior work has established that sex-specific VIC myofibroblast activation and drug sensitivity is better captured on hydrogel biomaterials over traditional TCPS^21,24^. As such, we investigated the effects of the eight selected inhibitors on a PEG hydrogel platform that mimics the stiffness of aortic valve tissue^26^. Using a custom-designed stamping tool, we developed a method to form hydrogels with flat topography suitable for cell culture on a 96-well glass-bottom plate (Fig. 2A-B, Supplementary Fig. 4). Using thiol-ene click chemistry, we developed soft and stiff hydrogel formulations that recapitulate the elastic modulus of healthy and diseased aortic valve tissue^50^ using PEG polymers of varying molecular weights and concentrations (Supplementary Fig. 5). The elastic moduli of the optimized soft and stiff hydrogel formulations were confirmed using oscillatory shear rheology (Fig. 2C).

**Fig. 2.**
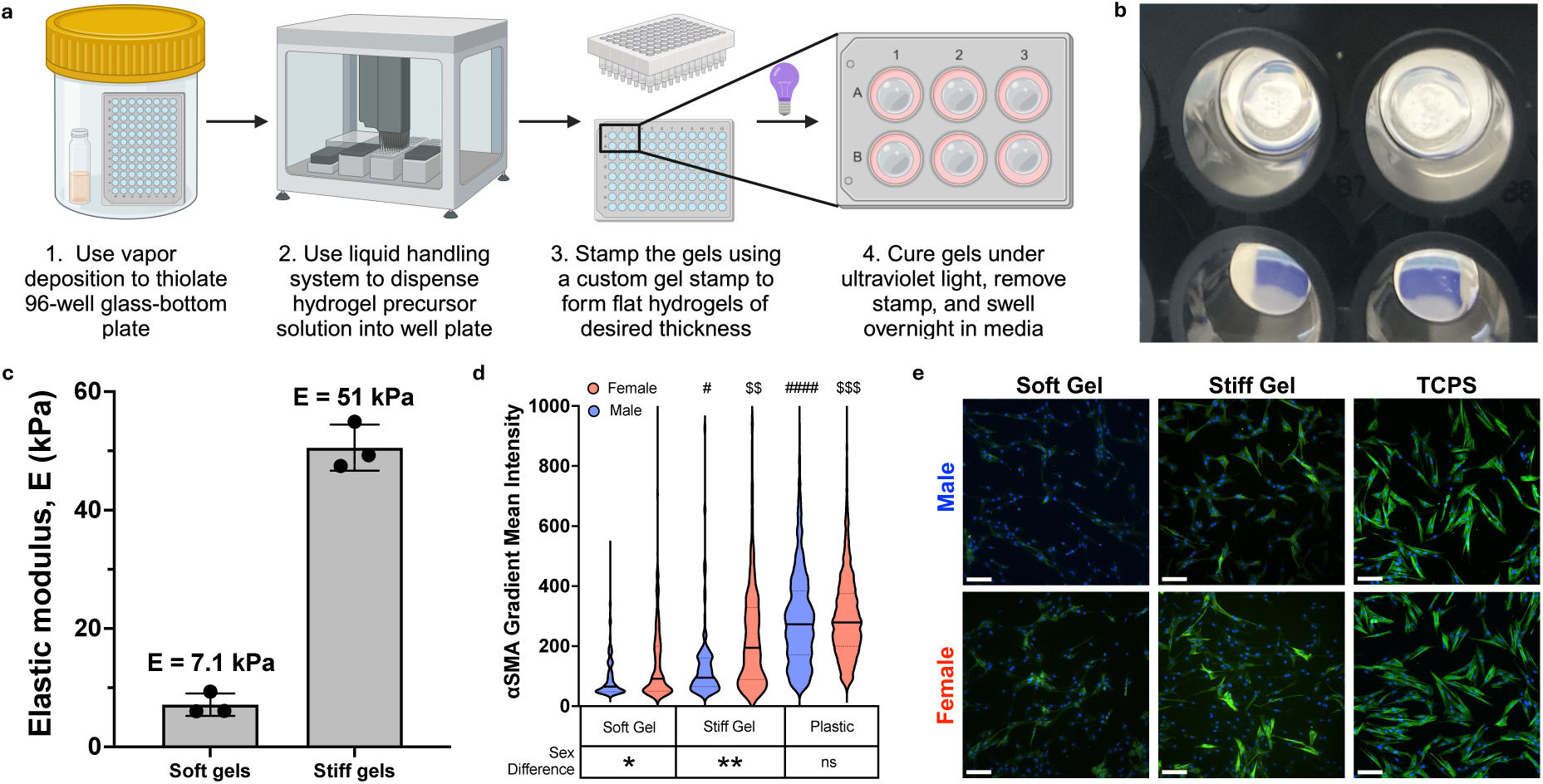
Developing a hydrogel platform for high-throughput drug screening. **a,** Schematic outlining the key steps to form hydrogels on a 96-well glass-bottom plate. **b,** Representative image of stiff hydrogels formed in a 96-well glass-bottom plate. **c,** Rheology of optimized soft and stiff gel formulations (n = 3 gels). **d,** Violin plot showing the distribution of αSMA gradient mean intensity of male and female VICs cultured on soft gels, stiff gels, or TCPS. A minimum of 500 cells were used per condition. **e,** Representative immunofluorescence images with αSMA stained in green and DAPI stained in blue. Scale bar = 100 μm. Statistical significance was determined by one-way ANOVA with Tukey posttests (P<0.0001) and Cohen’s d-test. # = Cohen’s d>0.2 and #### = Cohen’s d>1.2 relative to male soft gel. $$ = Cohen’s d>0.5 and $$$ = Cohen’s d>0.8 relative to female soft gel. * = Cohen’s d>0.2 and ** = Cohen’s d>0.5 between sexes.

We also confirmed that our new hydrogel platform reliably reproduces previously observed sex-specific VIC phenotypes. Aligning with previous findings, we observed a female-bias towards myofibroblast activation when VICs are cultured on soft or stiff hydrogels. Moreover, male and female VICs showed increased αSMA GMI as a function of increased cell culture substrate stiffness (Figs. 2D-E). Importantly, sex differences in myofibroblast activation are lost on TCPS due to the supraphysiologic stiffness (E∼1 GPa) inducing rampant myofibroblast activation equally in male and female VICs.

### Characterizing sex-specific anti-fibrotic drug dosing on stiff hydrogels

Using our high-throughput hydrogel platform, we next performed a drug screen with male and female VICs cultured on stiff hydrogels instead of TCPS (Figs. 3A-H). In contrast to the findings using TCPS, only four out of the eight anti-fibrotic drugs were more effective in male VICs (Fig. 3I). Moreover, comparing the EC_20_ values revealed that only Losartan elicited a sex-specific response by again reducing αSMA GMI more in male VICs relative to female VICs (Supplementary Fig. 6). Similar to the findings using TCPS, no inhibitors were significantly more effective in female VICs. Importantly, we confirmed the cytotoxicity for all inhibitors was minimal at low doses via the immunofluorescent intensity of cleaved caspase-3 (Supplementary Fig. 7).

**Fig. 3.**
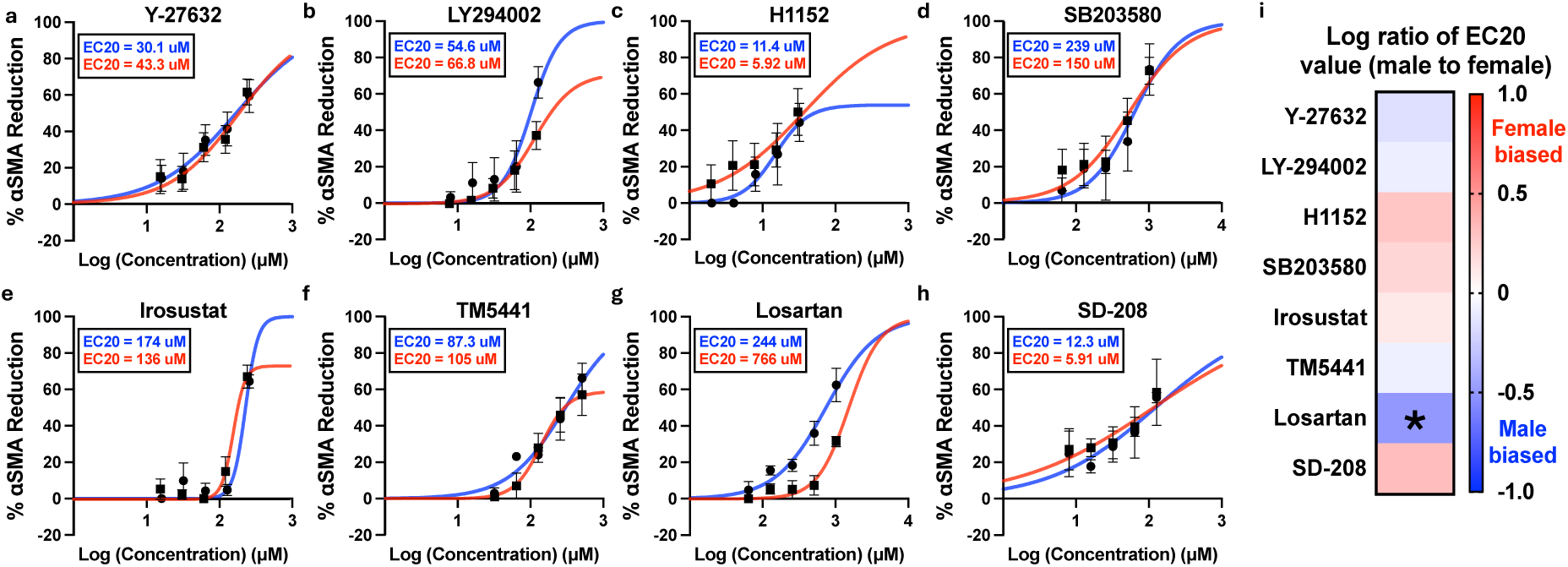
Drug response curves for eight anti-fibrotic drugs inhibiting male and female VIC myofibroblast activation on stiff gels. **a-h,** Percent αSMA reduction in male VICs (blue) and female VICs (red) cultured on stiff hydrogels with indicated doses of **a** Y-27632, **b** LY294002, **c** H1152, **d** SB203580, **e** Irosustat, **f** TM5441, **g** Losartan, and **h** SD-208. **i,** Heat map showing the log ratio of male EC_20_ to female EC_20_ for each inhibitor. Data is plotted as mean ± standard error of the mean (N = 3 biological replicates). The best fit line shown was generated using a nonlinear regression curve fit with GraphPad Prism. Statistical significance was determined by an unpaired two-tailed t-test with Welch’s correction and indicated as *=P<0.05.

We next investigated the combinatorial effects of anti-fibrotic drugs on male and female VICs. As an initial test, we probed the effects of combinations of a moderate and high dose of H1152 and LY294002 as these inhibitors were highly effective in reducing myofibroblast activation in male and female VICs individually. We found that H1152 and LY294002 interact to further reduce myofibroblast activation in male and female VICs, motivating further investigation into optimizing drug combinations with our eight drug library (Supplementary Fig. 8).

IDentif.AI analysis interrogates drug combinations in three concentration levels: Level 0 (L0), Level 1 (L1), and Level 2 (L2), which represent the absence of a drug and two clinically relevant concentrations, respectively. To establish doses of each drug that are non-cytotoxic for combinatorial analysis, we tested all eight inhibitors in a single combination at multiple doses with male and female VICs cultured on stiff hydrogels. Concentrations were selected based on dose response analysis data (EC value), cytotoxicity data (CC value), and pharmacokinetic data^45,46,51^ (10% of C_max_; maximum human serum from clinical trials). We found that combining all inhibitors at doses limited by EC_2_, CC_2_, or 10% of C_max_ led to substantial reductions in αSMA GMI in both male and female VICs with no apparent cytotoxicity (Supplementary Fig. 9). Thus, the L2 concentrations for each inhibitor were limited by EC_2_/CC_2_/10% of C_max_ while the L1 doses were limited by EC_1_/CC_1_/5% of C_max_ for male and female VICs (Supplementary Tables 2-3).

### IDentif.AI male VIC analysis pinpoints efficacious drug combinations based on interactions between Losartan and SD-208

We next harnessed the IDentif.AI platform^45,46,51^ to pinpoint combinatorial designs that may substantially increase the percent αSMA reduction in male VICs. First, male VICs cultured on stiff hydrogels were exposed to a set of 59 orthogonal array composite design^52^ (OACD) combinations at the L0, L1, and L2 concentrations for each inhibitor (Supplementary Tables 4 and 5). The monotherapies (L1/L2) of the inhibitors were also experimentally validated (Supplementary Table 6). The combinations along with monotherapies were correlated with their prospectively acquired percent αSMA reduction data via a second order quadratic series, which informed the parameter space of 6,561 (3^8^) combinations. A square transformation was applied to the percent αSMA reduction data to improve the goodness-of-fit of the model. The IDentif.AI analysis resulted in an adjusted R^2^ of 0.655, representing a well correlated relationship between the input combinations and their respective biological responses (Supplementary Table 7). No outliers were detected (Supplementary Fig. 10A) and all combinations were confirmed to be noncytotoxic (Supplementary Fig. 11A).

The male VIC IDentif.AI analysis yielded a ranked list of all combinations along with their predicted percent αSMA reduction. We prioritized three combinations from the top 2-drug, 3-drug, and 4-drug combinations (M1-M9) for subsequent experimental validation (Supplementary Table 8). Additionally, we included two multi-drug combinations that were predicted to be ineffective as negative controls (M10-M11). Eight out of the nine IDentif.AI-designed combinations that were predicted to be highly effective did indeed significantly reduce αSMA GMI in male VICs cultured on stiff hydrogels (Fig. 4A). Moreover, the two predicted ineffective drug combinations were confirmed to have a minimal impact on male VIC myofibroblast activation, and all combinations were well tolerated by the cells with minimal cytotoxicity (Supplementary Fig. 12A). Comparing the predicted and measured percent αSMA reduction of each combination revealed a significant correlation between model prediction and experimental output (Fig. 4B). As a final validation, three of the most effective drug combinations (M6, M7, and M8) and the least effective drug combination (M11) were tested with three biological replicates. The three predicted effective combinations significantly reduced αSMA gradient mean intensity in male VICs, while the predicted ineffective combination had no meaningful effect (Figs. 4C-D).

**Fig. 4.**
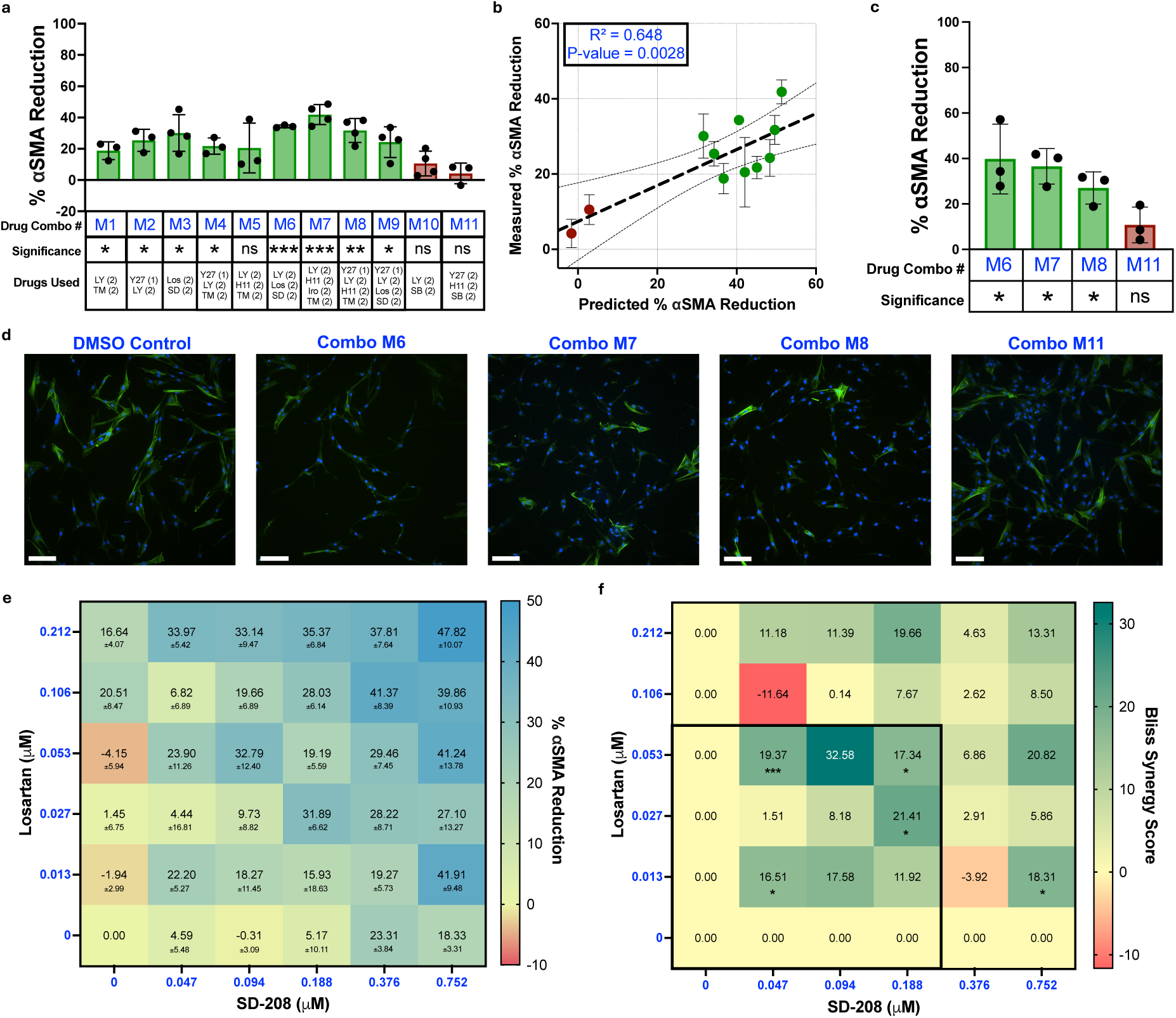
IDentif.AI analysis validation and drug synergy analysis for male VICs. **a,** Percent αSMA reduction in male VICs cultured with IDentif.AI-designed effective (green) and ineffective (red) drug combinations. (1) indicates L1 dose and (2) indicates L2 dose of that drug (n = 3-4). **b,** Correlation plot between the predicted vs measured percent αSMA reduction for IDentif.AI designed combinations. **c,** Validation of top three and bottom ranked IDentif.AI drug combinations (N = 3). **d,** Representative immunofluorescent images with αSMA stained in green and DAPI stained in blue. Scale bar = 100 μm. **e,** Interaction map of Losartan and SD-208 at various dose ratios and their respective percent αSMA reduction (N = 3). **f,** Synergy map of Losartan and SD-208 at various dose ratios (N = 3). The black box indicates the therapeutic dosing range (0 - L2) used for IDentif.AI analysis. Statistical significance was determined by a one sample t-test against a theoretical mean of zero and indicated as *=P<0.05, **=P<0.01, and ***=P<0.001. Bliss synergy scores of < -10, between -10 and 10, and > 10 represent antagonistic, additive, and synergistic drug interactions. Data is plotted as mean ± standard deviation. Abbreviations: H1152 (H11); Irosustat (Iro); Losartan (Los); LY294002 (LY); SB203580 (SB); SD-208 (SD); TM5441 (TM); Y-27632 (Y27). N = number of biological replicates. n = number of gels.

Furthermore, our IDentif.AI analysis pointed to interesting interactions between Losartan and SD-208. The interaction surface of Losartan and SD-208 indicated that, at L2 concentrations, the combination may exhibit interactions that further enhance therapeutic efficacy (Supplementary Fig. 13A). The significantly positive bilinear coefficient in the model representing the interaction between Losartan and SD-208 also suggests a strong interaction between the two drugs (Supplementary Table 7). Subsequently, we assessed 25 distinct dose ratios of Losartan and SD-208 via a 6×6 checkerboard assay (Fig. 4E). By comparing the percent αSMA reduction of the two drugs individually against when used in combination, we identified five dose ratios of Losartan and SD-208 that are synergistic as quantified by a significantly positive Bliss synergy score (Fig. 4F). Interestingly, four of the five synergistic dose ratios of Losartan and SD-208 were within the low-dose therapeutic range (< L2 concentration) used for IDentif.AI analysis.

### IDentif.AI female VIC analysis pinpoints efficacious drug combinations based on interactions between LY294002 and H1152

The IDentif.AI workflow was repeated to interrogate the drug-drug interaction space for female VICs cultured on stiff hydrogels. We followed the same workflow to screen the drug interaction space for female VICs (Supplementary Table 9). All drug combinations at female-specific doses were nontoxic and all inhibitors had similar therapeutic efficacy individually (Supplementary Fig. 11B and Supplementary Table 10). Similarly, IDentif.AI correlated the input OACD combinations and monotherapies with their measured percent αSMA reduction data via a second order quadratic series, resulting in an adjusted R^2^ of 0.651 (Supplementary Table 11). No transformation was applied to the percent αSMA reduction data, and no outliers were detected via a series of residual-based outlier analysis (Supplementary Fig. 10B).

Next, we tested three of the top 2-drug, 3-drug, and 4-drug combinations that were predicted to be highly effective in female VICs (F1-F9) along with three predicted ineffective multi-drug combinations (F10-F12) (Supplementary Table 12). We found that all three predicted ineffective drug combinations had no impact on female VIC myofibroblast activation, whereas six out of the nine predicted effective combinations induced meaningful reductions in αSMA GMI (Fig. 5A). All twelve drug combinations were confirmed to be noncytotoxic (Supplementary Fig. 12B). The female VIC IDentif.AI analysis resulted in a significant correlation with the *in vitro* data, suggesting high predictive accuracy similar to our male dataset (Fig. 5B). Three predicted effective IDentif.AI-designed combinations (F3, F4, and F7) and one predicted ineffective combination (F12) were further validated with additional biological replicates (Figs. 5C-D). Two out of the three predicted effective drug combinations were again confirmed to significantly reduce αSMA GMI, with the third trending toward significance (*P* = 0.067). Following previous results, the predicted ineffective drug combination had no significant impact on female VIC myofibroblast activation.

**Fig. 5.**
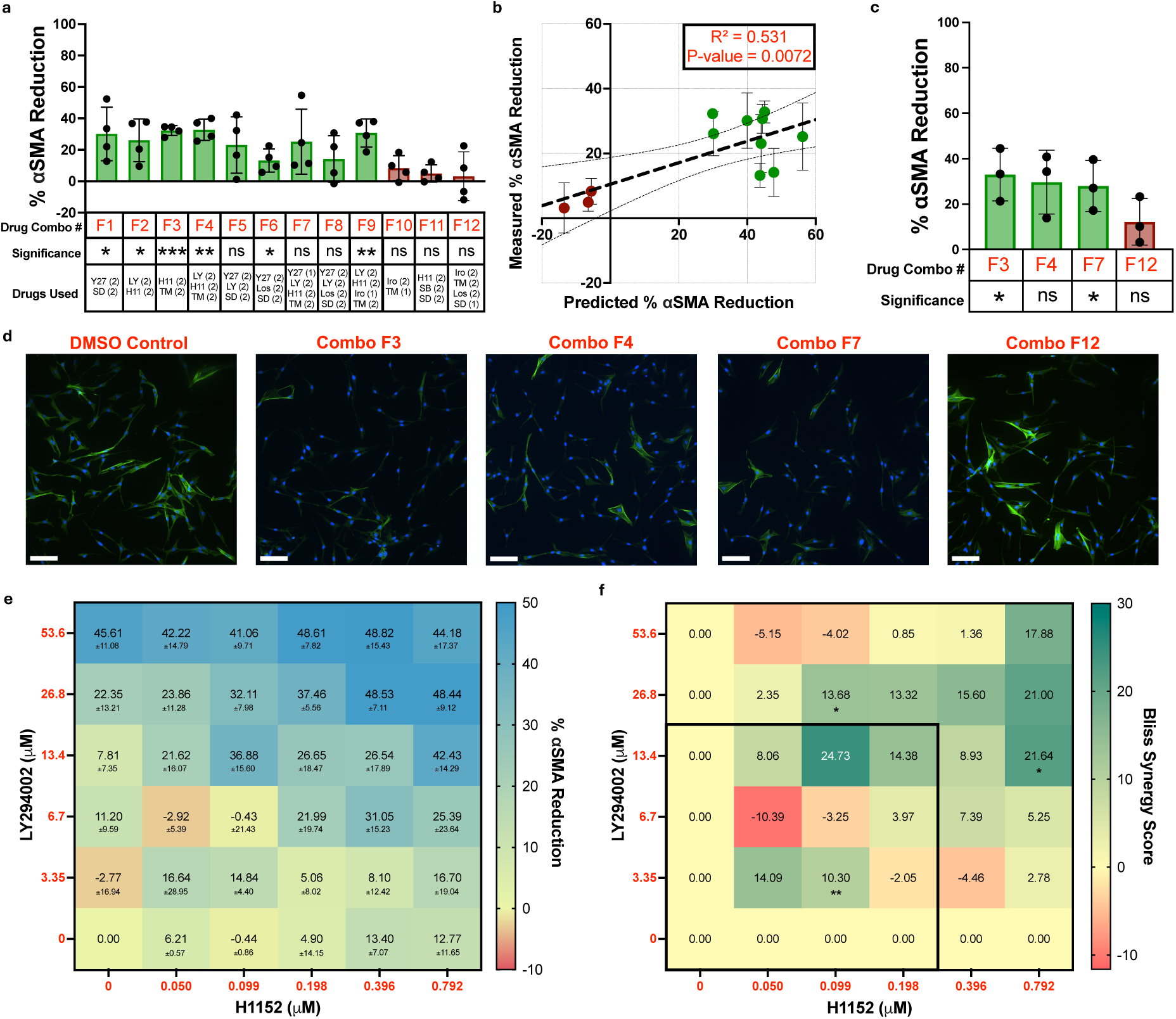
IDentif.AI analysis validation and drug synergy analysis for female VICs. **a,** Percent αSMA reduction in female VICs cultured with IDentif.AI-designed effective (green) and ineffective (red) drug combinations. (1) indicates L1 dose and (2) indicates L2 dose of that drug (n = 4). **b,** Correlation plot between the predicted vs measured percent αSMA reduction for IDentif.AI designed combinations. **c,** Validation of top three and bottom ranked IDentif.AI drug combinations (N = 3). **d,** Representative immunofluorescent images with αSMA stained in green and DAPI stained in blue. Scale bar = 100 μm. **e,** Interaction map of LY294002 and H1152 at various dose ratios and their respective percent αSMA reduction (N = 3). **f,** Synergy map of LY294002 and H1152 at various dose ratios (N=3). The black box indicates the therapeutic dosing range (0 - L2) used for IDentif.AI analysis. Statistical significance was determined by a one sample t-test against a theoretical mean of zero and indicated as *=P<0.05, **=P<0.01, and ***=P<0.001. Bliss synergy scores of < -10, between -10 and 10, and > 10 represent antagonistic, additive, and synergistic drug interactions. Data is plotted as mean ± standard deviation. Abbreviations: H1152 (H11); Irosustat (Iro); Losartan (Los); LY294002 (LY); SB203580 (SB); SD-208 (SD); TM5441 (TM); Y-27632 (Y27). N = number of biological replicates. n = number of gels.

In the female VIC analysis, IDentif.AI detected notable interactions between LY294002 and H1152 on stiff hydrogels, suggesting that there may be interactions when both drugs achieve L2 concentrations (Supplementary Fig. 13B). More importantly, a significantly positive bilinear term for the combination in the female VIC IDentif.AI analysis further suggests strong interactions between the two drugs (Supplementary Table 11). Interactions between LY294002 and H1152 were assessed via a 6×6 checkerboard assay (Fig. 5E). Out of the 25 different dose ratios, three emerged as synergistic as determined by a significantly positive Bliss synergy score (Fig. 5F). However, only one synergistic dose ratio of LY294002/H1152 was in the low-dose therapeutic range used for IDentif.AI analysis.

### IDentif.AI analysis pinpoints combinations and drug interactions that are sex-specific

We next assessed if the top ranked male and female drug combinations are sex-specific. We tested four of the top male combinations (M1, M4, M7, and M8) at male-optimized doses as well as four of the top female combinations (F1, F4, F7, and F8) at female-optimized doses in both male and female VICs cultured on stiff hydrogels. Except for M1, which was significantly more effective in male VICs, no drug combinations were sex-specific (Supplementary Fig. 14A). Grouping the drug combinations together as either male-optimized or female-optimized and subtracting the male and female percent αSMA reduction for each combination showed that both groups have an average percent αSMA reduction difference close to zero, indicating no meaningful sex difference between the groups (Supplementary Fig. 14B).

To identify candidate sex-biased combinations, we extracted all IDentif.AI-designed 2-drug, 3-drug, and 4-drug combinations along with their predicted percent αSMA reduction from both male and female VIC IDentif.AI analyses. For every combination, the predicted female percent αSMA reduction was subtracted from the predicted male percent αSMA reduction and ranked by the percent αSMA reduction difference (Supplementary Data 1). Combinations with a positive percent αSMA reduction difference were defined as male-biased and combinations with a negative percent reduction difference were defined as female-biased. The top male-biased combinations were most commonly composed of LY-294002, Irosustat, and TM5441, whereas the top female-biased combinations were most commonly composed of Y-27632, H1152, and SB203580 (Supplementary Fig. 15). We selected each of the top predicted male-biased (MB1-MB3) and female-biased (FB1-FB3) 2-drug, 3-drug, and 4-drug combinations for validation (Supplementary Table 13). Following IDentif.AI predictions, all male-biased drug combinations were more effective in male VICs and all female-biased drug combinations were more effective in female VICs (Fig. 6A). Comparing the predicted difference between male and female VIC percent αSMA reduction with the measured difference in percent αSMA reduction for each combination reveals a significant correlation (Fig. 6B). Additionally, further grouping the combinations as either male-biased or female-biased shows that, taken together, these classes of combinations are sex-specific (Fig. 6C).

**Fig. 6.**
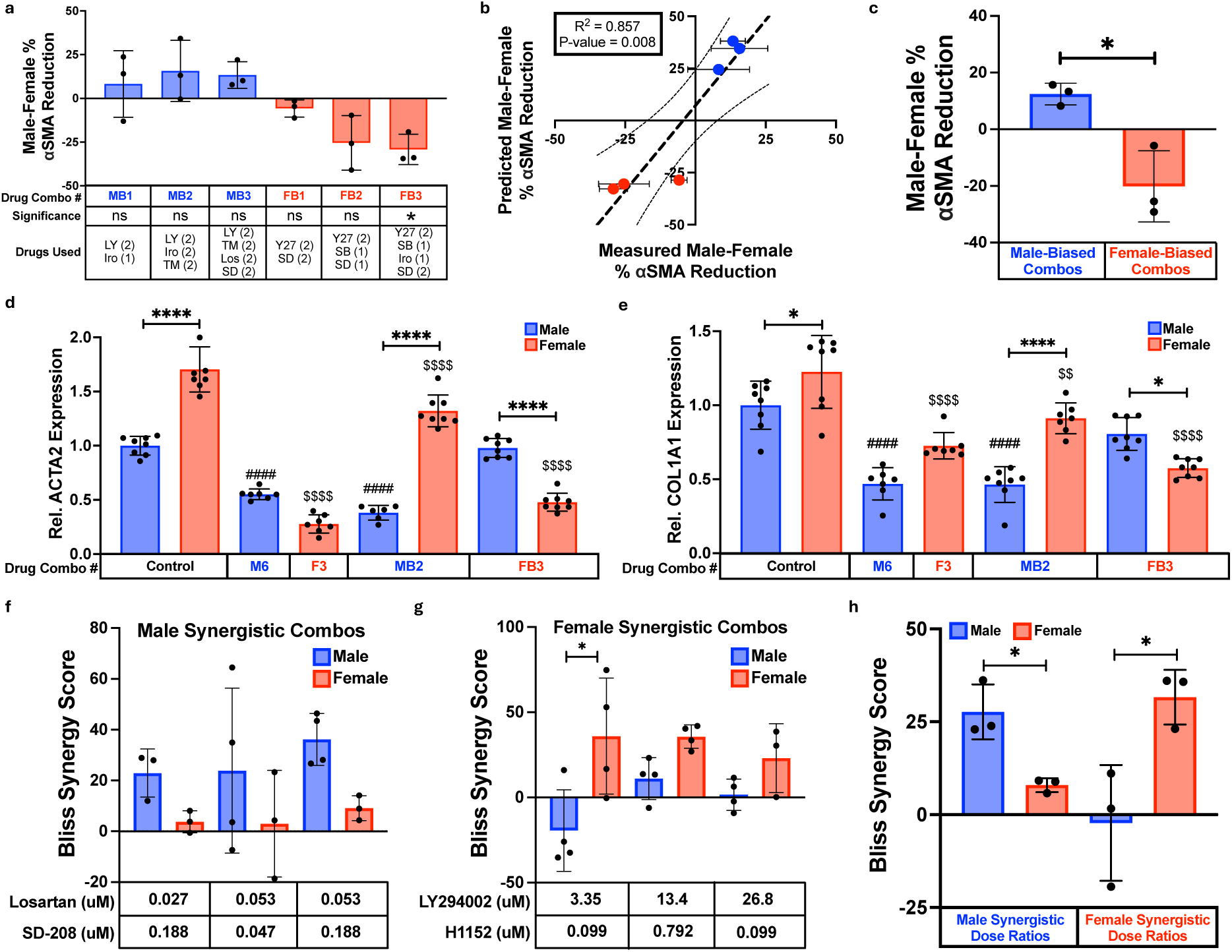
Comparing male-optimized and female-optimized drug combinations. **a,** Bar plot and **b** correlation plot of percent αSMA reduction difference (male-female) between male and female VICs cultured with top predicted male-biased and female-biased combinations. (1) indicates L1 dose and (2) indicates L2 dose of that drug (N = 3). **c,** Percent αSMA reduction difference between male and female VICs cultured with groups of predicted male-biased and female-biased combinations with three distinct drug combinations per group. **d,e,** Quantitative RT-qPCR of relative **d** ACTA2 and **e** COL1A1 gene expression in male and female VICs (n = 6-8). **f,** Bliss synergy scores for male and female VICs cultured with identified male synergistic dose ratios of Losartan and SD-208 on stiff hydrogels (n = 3-4). **g,** Bliss synergy scores for male and female VICs cultured with identified female synergistic dose ratios of LY294002 and H1152 on stiff hydrogels (n = 3-4). **h,** Bliss synergy scores for male and female VICs cultured with one of three male synergistic dose ratios of Losartan/SD-208 or one of three female synergistic dose ratios of LY294002/H1152. Statistical significance was determined for **a** by one sample t-test against a theoretical mean of zero, **c** and **h** by unpaired two-tailed t-test with Welch’s correction, and **d-g** by one-way ANOVA with Tukey posttests. For all graphs, significance between groups is indicated as *=P<0.05 and ****=P<0.0001 Significance relative to male control is indicated as ####=P<0.001 and significance relative to female control is indicated as $$=P<0.01 and $$$$=P<0.001. Bliss synergy scores of < -10, between -10 and 10, and > 10 represent antagonistic, additive, and synergistic drug interactions. Data is plotted as mean ± standard deviation. Abbreviations: Irosustat (Iro); Losartan (Los); LY294002 (LY); SB203580 (SB); SD-208 (SD); TM5441 (TM); Y-27632 (Y27). N = number of biological replicates. n = number of gels.

Next, we performed RT-qPCR to confirm our findings and to investigate how IDentif.AI-optimized drug combinations alter myofibroblast gene expression in response to the top male or female combination (M6 or F3) as well as the top sex-biased drug combinations (MB2 and FB3) on stiff hydrogels. Specifically, we assessed *ACTA2* and *COL1A1* expression as both genes are established phenotypic markers that are upregulated in myofibroblasts^53^. Aligning with our immunofluorescence findings, female VICs had significantly higher levels of *ACTA2* expression compared to male VICs at baseline (Fig. 6D). As expected, combinations M6 and MB2 lowered *ACTA2* expression in male VICs, whereas combination FB3 had no effect. In contrast, female VICs showed much larger reductions in *ACTA2* gene expression from combinations F3 and FB3 over combination MB2. Similar trends were observed when analyzing *COL1A1* gene expression (Fig. 6E). Male VICs had lower baseline *COL1A1* expression than female VICs and only had lowered *COL1A1* expression in response to combinations M3 and MB2, whereas female VICs experienced the largest reductions in *COL1A1* expression from combinations F3 and FB3.

We also wanted to determine if male and female synergistic combinations are sex-specific. We measured the percent αSMA reduction of three of the five identified synergistic dose ratios of Losartan/SD-208 as well as their individual effects with both male and female VICs to compute Bliss synergy scores (Fig. 6F). Similarly, we calculated Bliss synergy scores for the three identified synergistic dose ratios of LY294002/H1152 with male and female VICs (Fig. 6G). Comparing the effects of each dose ratio in male and female VICs shows that combinations of Losartan/SD-208 are more synergistic in male VICs and combinations of LY294002/H1152 are more synergistic in female VICs (Fig. 6H).

### IDentif.AI-designed combinations can reverse VIC myofibroblast activation and are most effective on soft and stiff hydrogels

Having confirmed the ability of IDentif.AI-designed combinations to prevent myofibroblast activation in male and female VICs, we next wanted to investigate if the optimized drug combinations were similarly effective in VICs activated to a myofibroblast prior to drug treatment. To test this, we altered our experimental setup to first culture VICs for either 2 days or 7 days to generate transiently or persistently activated VICs^54^, then add in drug combinations for 2 days (Fig. 7A). This approach determines if optimized combinations can *reverse* in addition to *prevent* myofibroblast activation in male and female VICs. We found that all three male optimized combinations (M6, M7, and M8) induced significant reductions in male VIC αSMA GMI after 2 days of culture (Fig. 7B). In contrast, one out of the three female optimized combinations (F6, F7, and F8) was significantly effective over the same time period in female VICs (Fig. 7C). In both male and female VICs, the predicted ineffective combos (M11 and F12) had no significant effect. We also tested the top male-biased (MB2) and top female-biased (FB3) combinations in male and female VICs after a 2 day culture (Fig. 7D). Surprisingly, we found that the sex-biased combinations were more sexually dimorphic at this longer timepoint with combination MB2 inducing a 36% greater αSMA reduction in male VICs and combination FB3 inducing a 59% greater αSMA reduction in female VICs.

**Fig. 7.**
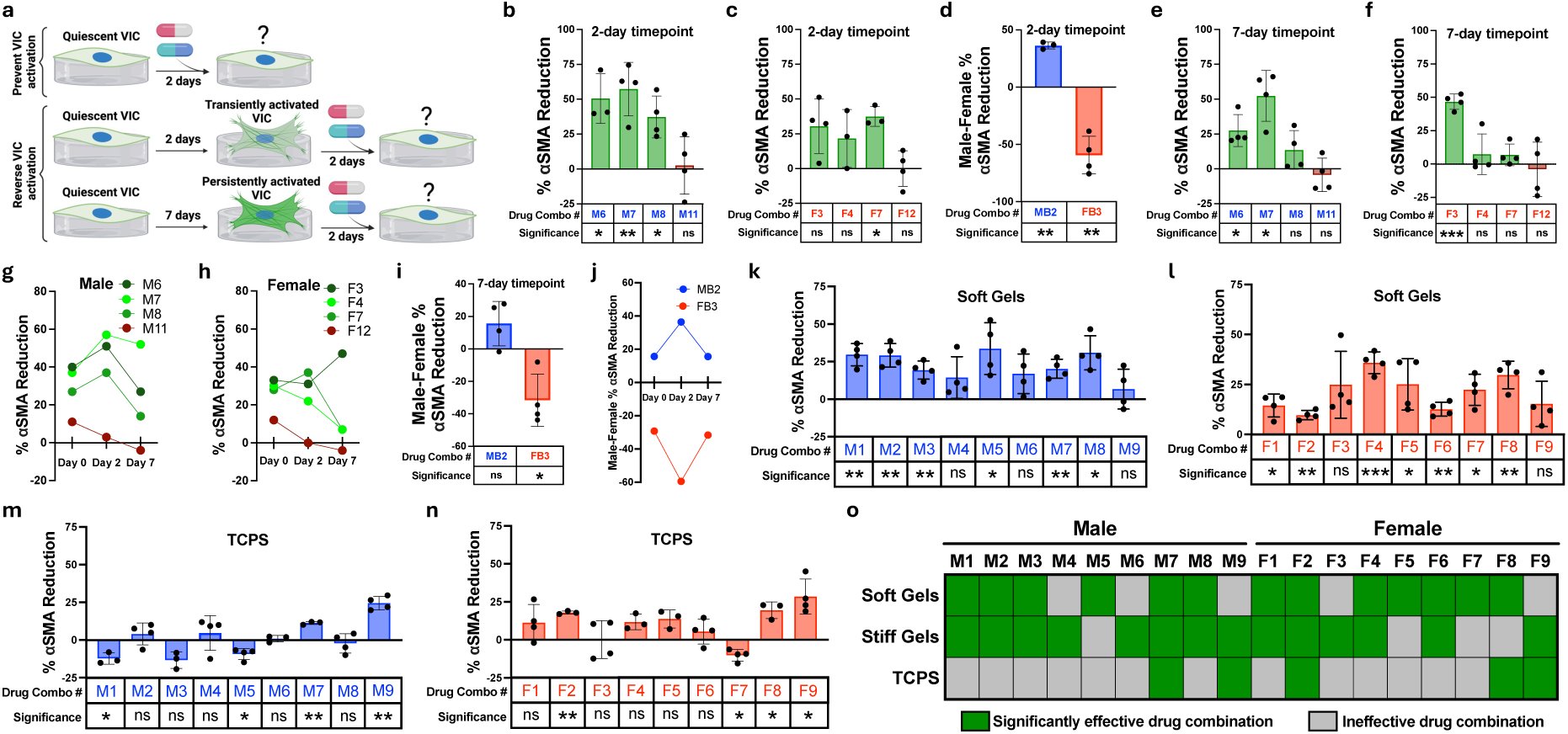
Quantifying the effects of cell culture timepoint and substrate stiffness on drug efficacy. **a,** Overview of experimental setup for assessing if drug combinations can reverse transiently and persistently activated VICs. **b-c,** Percent αSMA reduction in **b** male VICs and **c** female VICs after two days of culture on stiff hydrogels prior to exposure to drug treatment (n = 3-4 gels). **d,** Percent αSMA reduction difference between male and female VICs after two days of culture on stiff hydrogels prior to exposure to drug treatment (n = 3-4 gels). **e,f,** Percent αSMA reduction in **e** male VICs and **f** female VICs after seven days of culture on stiff hydrogels prior to exposure to drug treatment (n = 4 gels). **g-h** Percent αSMA reduction in **g** male VICs and **h** female VICs after zero, two, or seven days of culture on stiff hydrogels prior to exposure to drug treatment. **i,** Percent αSMA reduction difference between male and female VICs after seven days of culture on stiff hydrogels prior to exposure to drug treatment (n = 4 gels). **j,** Percent αSMA reduction difference between male and female VICs after zero, two, or seven days of culture on stiff hydrogels prior to exposure to drug treatment. **k,l,** Percent αSMA reduction in **k** male or **l** female VICs cultured with the top drug combinations on soft hydrogels (n = 3-4 gels). **m,n,** Percent αSMA reduction in **m** male or **n** female VICs cultured with the top drug combinations on TCPS (n = 3-4 wells). **o,** Summary heatmap showing which optimized male and female drug combinations are significantly effective on soft hydrogels, stiff hydrogels, and TCPS. Data is plotted as mean ± standard deviation. Statistical significance was determined by a one sample t-test against a theoretical mean of zero and indicated as *=P<0.05, **=P<0.01 and ***=P<0.001.

To test the effects of drug combinations in persistently activated myofibroblasts, we extended the VIC culture period to 7 days prior to adding in drug combinations. We observed combinations M6 and M7 maintained a significant therapeutic effect, whereas combination M11 was rendered ineffective in male VICs (Fig. 7E). In female VICs, combination F3 emerged as the only combination to induce a significant reduction in αSMA GMI in female VICs after 7 days of culture (Fig. 7F). Moreover, combination F3 was the only combination that was more effective in reversing myofibroblast activation after 7 days of culture compared to 2 days of culture in male or female VICs (Figs. 7G-H). After the 7 day culture, combination MB2 trended toward being more effective in male VICs (*P* = 0.11) and combination FB3 was still more effective in female VICs (Fig. 7I). However, both combinations were less sex-specific after a 7 day culture relative to a 2 day culture as indicated by percent αSMA reduction differences closer to zero (Fig. 7J).

Lastly, we tested the ability of IDentif.AI-designed combinations to prevent myofibroblast activation in VICs cultured on other substrates beyond stiff hydrogels. Thus, we cultured male VICs on soft hydrogels and added in each of the nine optimized male combinations (M1-M9). Six male-optimized combinations were effective in preventing male VIC myofibroblast activation (Fig. 7K). Similarly, in female VICs, seven out of the nine female-optimized combinations (F1-F9) generated significant reductions in αSMA GMI on soft hydrogels (Fig. 7L). Repeating the same experiments with TCPS led to significantly different results, with only two male-optimized combinations and three female-optimized combinations being effective in male and female VICs, respectively (Figs. 7M-N). Interestingly, combination M7 in male VICs and combination F2 in female VICs were significantly effective across soft hydrogels, stiff hydrogels, and TCPS (Fig. 7O).

## Discussion

Collectively, our work identified and validated sex-specific drug combinations to inhibit VIC myofibroblast activation using our high-throughput hydrogel cell culture platform. Hydrogel precision biomaterials were optimized to mimic the mechanical properties of healthy and diseased valvular tissue, enabling us to recapitulate sex-specific VIC myofibroblast activation *in vitro*^55,56^ (Fig. 2). Multiple reports have shown that hydrogels reproduce phenotypes in a variety of cell types for improved drug screening, including cardiomyocytes^57^, glioblastoma cells^58^, and hepatocytes^59^. In our study, we utilized hydrogel biomaterials to evaluate the efficacy of anti-fibrotic drugs in male and female VICs instead of materials with supraphysiologic stiffness such as TCPS (Figs. 1 and 3). We found that optimized drug combinations were significantly less effective when VICs were cultured on TCPS over soft and stiff hydrogels, indicating an interplay between cell culture substrate stiffness and drug efficacy (Fig. 7). This is likely due to the abnormal upregulation of cytoskeletal and contractility associated genes in VICs cultured on TCPS^60^ overriding the anti-fibrotic effects of optimized drug combinations. Interestingly, combinations M7 and F2 were effective across all cell culture substrates tested, and both contained LY294002 and H1152 (Fig. 7). The ability of H1152 to target VIC mechanosensing through inhibiting ROCK signaling combined with LY294002 inhibiting the PI3K biochemical signaling pathway offers an explanation for the efficacy of M7 and F2 across multiple cell culture substrates. However, other drug combinations that also contained LY294002 and H1152 (M5, M8, F4, F7, and F9) were not effective across all substrates, suggesting that multi-drug interactions may also play a role in the robustness of drug combination efficacy (Fig. 7).

Our hydrogel platform also enabled the assessment of drug combinations to reverse transient and persistent VIC myofibroblast activation. In healthy valve tissue, VICs become transiently activated to myofibroblasts to perform routine matrix remodeling then return to a quiescent state after the remodeling process is complete. However, after chronic valve injury, VICs can transition to persistently activated myofibroblasts with an irreversible phenotype and remain activated throughout several decades of AVS progression^61^. Prior work has been able to recapitulate transient and persistent valve myofibroblast activation found in diseased tissues via time-dependent mechanical “dosing” of VICs on stiff hydrogels *in vitro*^54^. Using previously established parameters, we found that subsets of optimized drug combinations that prevent VIC myofibroblast activation were also able to reverse transiently and persistently activated VICs (Fig. 7). Given our drug combinations were able to reduce persistent myofibroblast activation, we suggest that the drug combinations identified in this paper warrant further investigation using humanized *in vitro* and/or *in vivo* models of valve disease.

Our work also demonstrates how AI-derived platforms may be leveraged to accelerate sex-specific precision medicine. Our study utilized IDentif.AI to determine sex-specific drug combinations that effectively inhibit myofibroblast activation in male and female VICs. Our approach is one example of a larger trend toward implementing computational tools for predicting effective precision drug treatments^62^. For example, one recent study incorporated estrogen signaling into a network model for cardiac fibrosis to develop sex-specific treatment recommendations^63^. Other researchers have leveraged differential gene expression in cardiac tissue to build *in silico* drug models that predict the sex-specific effects of drugs targeting the heart^64^. However, most sex-specific drug prediction models are based on large -omics datasets. In contrast, IDentif.AI relies solely on small, prospectively generated datasets, enabling the further customization of drug combinations in future studies (Supplementary Tables 5 and 9). We suggest that IDentif.AI may serve as an approach to rapidly determine targeted drug combinations for AVS and other diseases that require precision medicine approaches and have a paucity of data available to develop predictive drug models.

Our data also provides a framework for rapidly optimizing drug combinations and dosing for clinically approved and/or experimental drugs in preclinical development. Among the eight actionable drugs we tested that are known to inhibit porcine valve myofibroblasts^29^, only Losartan, a potent inhibitor of the angiotensin II type 1 receptor used to treat hypertension, is approved for medical use. Losartan has been shown to reduce fibrosis in patients with nonobstructive hypertrophic cardiomyopathy^65^, demonstrating its clinical relevance in addressing cardiovascular disease. Moreover, Losartan was the only inhibitor to elicit sex-specific responses on TCPS and stiff hydrogels by inducing greater reductions in αSMA GMI in male VICs (Figs. 1 and 3). Beyond individual drug effects, significant sex differences in drug interactions were observed with female VICs being more responsive to LY294002/H1152 and male VICs being more responsive to Losartan/SD-208 (Fig. 6). In our study, Losartan was tested in clinically relevant concentrations and therefore, synergistic interactions observed between Losartan and SD-208 in male VICs are clinically relevant and may assist in repurposing already approved drug candidates for AVS. Furthermore, we anticipate the IDentif.AI workflow may accelerate the development and approval process of investigational or preclinical drugs (e.g. SD-208, LY294002, and H1152) and provide the backbone of combinatorial designs for AVS-specific drug development.

Collectively, we anticipate IDentif.AI and other AI-based tools will be critical for determining precision drug combinations and dosage parameters for AVS patients. For example, in the Simvastatin and Ezetimibe in Aortic Stenosis (SEAS) trial (NCT00092677), Simvastatin and Ezetimibe were paired in combination to evaluate the composite of major cardiovascular events including aortic valve replacement^12^. However, the one dose ratio tested did not reduce the number of cardiovascular events when compared to the placebo arm. In this study, drug combinations are sex-specific and may result in unforeseen interactions at different dose ratios (Figs. 4 and 5). Our work also suggests that the rapid evaluation of drug combinations and dosing parameters using IDentif.AI may inform future clinical trial design^42,43^. Even though our current work was limited to evaluating porcine myofibroblasts, we anticipate future humanized *in vitro* models using human derived VICs^22,66^, human serum^25,67^, and three-dimensional cell culture^68^ will rapidly converge on translationally relevant and sex-specific drug dosing parameters. In sum, continuous optimization of experimental models may accentuate important information and resources that accelerate targeted AVS drug development.

## Methods

### 96-well hydrogel fabrication

Glass-bottom 96-well plates (Cellvis) were placed in an autoclave jar with a scintillation vial containing 100 μL of mercaptopropyltrimethoxysilane (MPTS, Sigma-Aldrich). The autoclave jar was then lightly sealed and placed inside an oven at 60°C for a minimum of 6 hours. The autoclave jar was opened in a chemical fume hood and the vapor deposited well plate was stored for up to two hours until ready for use. 8-arm 20 kDa polyethylene glycol-norbornene (PEG-Nb) was formed as outlined previously^69^. A 7% wt/vol PEG-Nb hydrogel precursor solution was formed by combining phosphate buffered saline (PBS), 20 kDa PEG-Nb at a 0.98:1 thiol-to-ene ratio, 5 kDa PEG-dithiol (Jenkem), CRGDS cell binding motif (Bachem), and lithium phenyl-2,4,6-trimethyl-benzoylphosphinate (Fisher Scientific). An EPMotion 5073l automated liquid handling system (Eppendorf) was used to pipet 7 μL of the hydrogel precursor solution into the center of each well of the vapor deposited 96-well glass-bottom plate. The well plate with the hydrogel precursor solution was then stamped using a custom 96-well plate stamp, inverted, and cured under UV light at 10 mW/cm^2^ for 3 minutes to induce gelation. The stamp was carefully removed by hand and the newly formed gels were sterilized with a solution of 5% isopropyl alcohol (Thermo Fisher) in PBS for 20 minutes. Gels were then washed twice with PBS and stored in VIC media composed of Media199 (Life Technologies), 1% fetal bovine serum (FBS, LifeTechnologies), 1 μg mL amphotericin B (Thermo Fisher), 50 U/mL penicillin (Sigma), and 50 μg/mL streptomycin (Sigma). Finally, the hydrogels were placed in an incubator at 37°C with 5% carbon dioxide for up to 24 hours until experimental use.

### Rheology

30 μL of hydrogel precursor solution was photopolymerized *in situ* using an Omnicure at 10 mW/cm^2^ for three minutes. A DHR3 rheometer (TA Instruments) and an 8 mm diameter parallel plate geometry were used to perform oscillatory shear rheology with an amplitude of 1% and frequency of 1 Hz. For each replicate, storage (G’) and loss modulus (G’’) measurements were taken for 100 seconds prior to photopolymerization, 150 seconds during photopolymerization, and 50 seconds after photopolymerization. Young’s modulus (E) measurements were then determined from the storage modulus using the equation E=2G’(1+ ***v***) where ***v*** is Poisson’s ratio which is assumed to be 0.5 as G’>>>G’’.

### VIC isolation

VICs were isolated directly from 6-8-month-old male and female adult pigs (Midwest Research Swine) following methods described previously^70^. Briefly, aortic valve leaflets were dissected, sex-separated, and immediately immersed into a solution of Earle’s Balanced Salt Solution (EBSS, Sigma-Aldrich) supplemented with 50 U/mL of penicillin (Thermo Fisher), 1 μg/mL amphotericin B (Thermo Fisher), and 50 μg/mL streptomycin (Thermo Fisher). Leaflets from a minimum of 4 sex-matched hearts were pooled together for each isolation, representing one biological replicate. After dissection, valve leaflets were transferred to a fresh solution of EBSS supplemented with 250 units type II collagenase (Worthington) per mL and shaken for 30 minutes inside an incubator at 37°C with 5% carbon dioxide. Leaflets were then vortexed on high for 30 minutes, placed in a fresh collagenase solution, and returned to an incubator for 60 minutes while being shaken. The digested valve tissue was again vortexed on high for two minutes, then pressed through a 100 μm cell strainer. VICs were then isolated through a 10-minute centrifugation at 0.2g to collect a cell pellet, which was resuspended with 10 mL of growth medium comprised of Media 199 (Life Technologies), 15% fetal bovine serum (FBS, Life Technologies), 1 μg/mL amphotericin B, 50 U/mL penicillin, and 50 μg/mL streptomycin. Next, VICs were seeded on T75 Falcon tissue culture treated flasks (Fisher) at 37°C and 5% carbon dioxide for expansion. Media changes were performed every 24 to 48 hours until VICs reached ∼80% confluency. Lastly, VICs were frozen down overnight in 1 mL aliquots comprised of 500 μL of FBS, 450 μL of 15% VIC media, and 50 μL of DMSO in freezing containers placed in a -80°C freezer. The next day, cell aliquots were frozen down in liquid nitrogen for long-term storage.

### VIC culture

For experiments, a frozen aliquot of passage 1 male or female VICs were thawed, resuspended in 9 mL of 15% VIC growth media and centrifuged at 0.2g for 5 minutes. After aspirating off the spent media, VICs were resuspended in 5 mL of fresh 15% FBS media and seeded on T25 Falcon tissue culture treated flasks (Fisher). Media changes were performed every 24-48 hours until cells reached ∼80% confluency. VICs were collected by aspirating off spent media, washing with phosphate buffered saline (PBS, Sigma-Aldrich) for 1 minute, then incubating with 1x trypsin (Life Technologies) for 3 minutes. Trypsin was neutralized with an equal volume of 15% FBS growth media and centrifuged for 5 minutes at 0.2g. The media solution was then aspirated and VICs were resuspended with 1 mL of 1% FBS media. VICs were counted with an automated hemocytometer to calculate the volume of cell solution to add to each well to seed at 6400 cells/well (20,000 cells/cm^2^) for TCPS and hydrogel experiments. For all experiments, VICs were seeded with 1% FBS media containing the specified drug doses and incubated at 37°C with 5% carbon dioxide with media changes every 48-72 hours if needed.

### Drug preparation and dosing

Lyophilized drug powders were resuspended with sterile filtered deionized water (H-1152 (Tocris) and Y-27632 (Tocris)) or sterile DMSO (LY-294002 (Cell Signaling Technology), SB203580 (Selleck Chemicals), Irosustat (Selleck Chemicals), TM-5441 (MedChemExpress), SD-208 (Sigma), Losartan (Selleck Chemicals), Ibrutinib (Selleck Chemicals), Bosentan (Sigma), and KDOAM-25 (MedChemExpress)). Drug stock solutions were discarded after two thaw cycles or after ten months of storage. For all experiments, drug dilutions were calculated and added to VIC media to ensure total DMSO levels remained below 1% for cytotoxicity and αSMA reduction curve fitting experiments and below 0.3% for drug combination experiments.

### Immunostaining

After culture on hydrogels or TCPS, the spent drug-containing media was removed and VICs were immersed in a 4% paraformaldehyde solution in PBS for 20 minutes then permeabilized for one hour with 0.1% Triton-X-100 (Fisher Scientific) in PBS. Cells were then blocked with 5% bovine serum albumin in PBS for 1 hour at room temperature or over the weekend at 4°C. Immunostaining was performed with a mouse anti-αSMA primary antibody (Abcam, 1:300) and a rabbit anti-cleaved Caspase 3 primary antibody (Abcam, 1:250 dilution) in blocking solution for one hour at room temperature. Next, VICs were washed with PBS with 0.05% Tween20 (Sigma) for 10 minutes and cultured in the dark for one hour with a secondary staining solution of PBS with goat anti-mouse Alexa Fluor 488 (Life Technologies, 1:300 dilution), goat anti-rabbit Alexa Fluor 647 (Life Technologies, 1:300 dilution), 4′-6-diamidino-2-phenylindole (Life Technologies, 1:500 dilution) and HCS Cell Mask Deep Orange (Life Technologies, 1:5000 dilution). Finally, VICs were washed once with PBS then stored in PBS at 4°C until imaging. Gel samples were imaged *in situ* using a Nikon Eclipse Ti2-E and analyzed using an automated MATLAB code to quantify the αSMA gradient mean intensities and caspase-3 intensities normalized to cell mask for each cell. A minimum of 100 cells were used per gel with multiple gels used per condition.

### αSMA reduction and cytotoxicity curve fitting

Curve fitting for VIC percent αSMA reduction and cytotoxicity was performed using GraphPad Prism 10 (GraphPad Software Inc). For each inhibitor, the nonlinear regression tool in Prism was used to estimate the response against the agonist by fitting the top, bottom, EC_50_/CC_50_, and hillslope to each data set. After 2 days of culture, the percent αSMA reduction was calculated using the average αSMA gradient mean intensities of the control relative to the drug condition using Eq. (1):

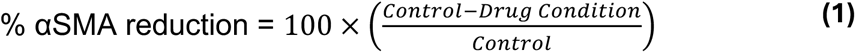

Percent cytotoxicity was determined by designating each cell as alive or dead using caspase-3 intensity normalized to cell mask intensity, which was thresholded using a positive and negative control for cell death (10 nM Staurosporine (Cell Signaling Technologies) or 1% FBS media). The bottom value for every curve fit was set to 0 and the top value was constrained to a maximum of 100 to indicate the maximum possible effect of the drug. The hillslope for each curve was limited to a maximum value of 5 to ensure a smooth curve fit and the EC_50_/CC_50_ variable was estimated without restrictions. Absolute EC and CC values for % αSMA reduction and percent cytotoxicity were quantified using the top, bottom, EC_50_/CC_50_, and hillslope values estimated from the curve fits generated in GraphPad Prism (GraphPad Software Inc).

### Harnessing IDentif.AI to optimize effective combinations for male and female VICs

The initial dose response analysis of all eight inhibitors along with pharmacokinetics data were considered to determine low-dose, non-cytotoxic, and clinically relevant L0, L1, and L2 concentrations for IDentif.AI analysis. Specifically, L0 corresponded to no drug (0 μM), L1 was obtained from EC_1_, CC_1_, and 5% of C_max_ for each drug, and L2 was obtained from EC_2_, CC_2_, and 10% of C_max_ (considered as an achievable concentration at the target tissue) for each drug per sex^44,46,47^ (Supplementary Tables 2 and 3). To interrogate the drug interaction space using IDentif.AI, a curated list of 59 combinations according to a resolution IV OACD^52^ was prepared and validated in male and female VICs. This design consists of 32 two-level fractional factorial combinations and 27 three-level orthogonal array combinations (Supplementary Table 4), which represent the minimum number of combinations needed to effectively screen the drug interaction space. The experimentally derived *in vitro* data from the 59 OACD combinations and monotherapies (L1/L2) were used to interrogate the drug-drug interaction space via a second order quadratic series derived using stepwise regression^44,46,51^ (MATLAB 2020R, MathWorks Inc). In addition, Box-Cox transformation was applied to the percent αSMA reduction data of OACD combinations and monotherapies (L1/L2) to determine potential transformations that may improve the residual distribution and goodness-of-fit of the model (e.g. adjusted R^2^). Outliers were determined via a series of residual-based outlier analysis. Utilizing the second order quadratic series, IDentif.AI provided a ranked list of all 6,561 combinations (3^8^; three concentration levels for eight drugs) and their predicted percent αSMA reduction.

### Drug synergy analysis

The most promising two drug interactions for male and female VICs based on the IDentif.AI analyses were further evaluated through a 6×6 checkerboard assay. Each drug was tested at six doses ranging from 0 μM up to four times the L2 dose with two-fold dilutions between each dose. The measured percent αSMA reduction data was used to generate an interaction map for each combination. Subsequently, the percent αSMA reduction data was uploaded to SynergyFinder to assess and quantify the interactions at each dose ratio for each drug combination using the Bliss independence model^44,46,51,71^. The resulting Bliss synergy scores for each dose ratio were used to generate synergy maps for the selected combinations. Bliss synergy scores of < -10, between -10 and 10, and > 10 represent antagonistic, additive, and synergistic drug interactions, redpectively^71^.

### Identifying sex-biased combinations

All IDentif.AI-pinpointed 2-drug, 3-drug, and 4-drug combinations along with the associated predicted percent αSMA reduction in male and female VICs were exported to Excel in separate tabs based on the number of drugs in each combination (Supplementary Data 1). In the male VIC IDentif.AI analysis, a square transformation was applied to the percent αSMA reduction data of the 59 OACD combinations. After the analysis, a square root transformation (inverse of square transformation) was performed to the predicted percent αSMA reduction data of all 6,561 combinations in order to restore the data to its original form. As a result of the transformations, a small portion of the low-ranked combinations containing negative values resulted in imaginary numbers, which were assumed to have a percent αSMA reduction of zero or no efficacy. All drug combinations were aligned between the male and female IDentif.AI analyses and the predicted percent αSMA reduction in female VICs was subtracted from the predicted percent αSMA reduction in male VICs. Drug combinations were then ranked based on the predicted percent αSMA reduction difference between male and female VICs from high to low, where positive numbers indicate a predicted male-bias and negative numbers indicate a predicted female-bias.

### RT-qPCR

For RT-qPCR, stiff hydrogels were formed directly on 25 mm circular glass coverslips that were thiolated via overnight vapor deposition with a vial of 100 μL of MPTS. After following the same sterilization steps outlined earlier, VICs were seeded (180,000 cells/gel) onto the hydrogels and cultured for 48 hours. VICs were then isolated from the hydrogels by inverting the gel into lysis buffer (Qiagen RNEasy Micro Kit) for two minutes and washing the coverslip repeatedly with 70% ethanol. Following the Qiagen RNEasy Micro Kit instructions, RNA was isolated and assessed for purity via a NanoDrop 2000 spectrophotometer (Thermo Fisher). All RNA had A260/A280 ratios between 1.8 and 2.1. The isolated RNA was immediately converted into cDNA following an iScript Synthesis kit (Bio-Rad) and stored at -20°C. Gene expression for *ACTA2* and *COL1A1* was assessed using iQ SYBR Green Supermix and a CFX384 iCycler (Bio-Rad). *ACTA2* and *COL1A1* were normalized to *RPL30* expression. All primer sequences can be found in Supplementary Table 14.

### Statistical Analysis

Unless otherwise stated, data are presented in mean ± standard deviation. N = number of biological replicates. n = number of wells or hydrogels. Unless otherwise stated, statistical significance was defined as a *P*-value less than 0.05 and calculated for comparisons between multiple groups via one-way ANOVA with Tukey posttests in GraphPad Prism (GraphPad Inc). Comparisons between two groups were calculated by unpaired two-sided *t*-test with Welch’s correction and one-way comparisons were calculated by one-way *t*-test against an assumed mean of zero in GraphPad Prism (GraphPad Inc). IDentif.AI analysis was performed using stepwise regression analysis in MATLAB (MATLAB 2020R, MathWorks Inc), and the statistics of IDentif.AI analysis were based on the sum-of-squares *F-*test.

### Data availability

All data can be found in the manuscript, supplementary information, and supplementary data.

### Code availability

A sample IDentif.AI code (MATLAB compatible) is provided in the supplementary information.

## Acknowledgements

B.V. acknowledges funding from the National Science Foundation Graduate Research Fellowship Program (NSF-GRFP DGE-2038238). Any opinions, findings, and conclusions or recommendations expressed in this material are those of the authors and do not necessarily reflect the views of the National Science Foundation. D.H. gratefully acknowledges funding from the Institute for Digital Medicine Translational Research Programme (grant number A-0001319-00-00) at the Yong Loo Lin School of Medicine, The N.1 Institute for Health, and the College of Design and Engineering at the National University of Singapore. B.A.A. acknowledges support from the National Institutes of Health (R00 HL148542), the NIH Director’s New Innovator Award (DP2 HL173948), the Chan Zuckerberg Initiative Science Diversity Leadership Award, and the American Heart Association (942253).

## Author contributions

B.J.V. and B.A.A. conceived and supervised the study. B.J.V. and M.C. performed all *in vitro* experiments. B.J.V., P.W., and M.C. analyzed data and performed statistical analyses. E.K.-H.C. and D.H. supervised the IDentif.AI workflow. P.G. assisted in developing the image analysis code. B.J.V., P.W, D.H., and B.A.A. wrote and edited the manuscript. All authors approved the manuscript.

## Materials and correspondence

All correspondence should be addressed to Brian Aguado, Dean Ho, and Edward Chow.

