## Supplementary information for "Determining sex differences in drug combinations targeting aortic valve myofibroblast activation using an artificial intelligence derived platform"

This Supplementary Information includes:

- Supplementary Tables 1-14
- Supplementary Figs. 1-15
- Example Code

**Supplementary Table 1 | Eleven inhibitors and corresponding therapeutic targets to reduce myofibroblast activation.**

| Drug | Therapeutic Target |
|---|---|
| Y-27632 | Rho-kinase (ROCK) 1 and ROCK 2 |
| LY294002 | Phosphoinositide 3-kinase α (PI3Kα), PI3Kδ, PI3Kβ |
| H1152 | ROCK2 |
| SB203580 | p38 mitogen-activated protein kinase |
| Irosustat | Steroid sulfatase |
| TM5441 | Plasminogen activator inhibitor 1 |
| Losartan | Angiotensin II type 1 receptor |
| SD-208 | Transforming growth factor beta receptor type 1 |
| Ibrutinib | Bruton’s tyrosine kinase and BMX non-receptor tyrosine kinase |
| KDOAM25 | Lysine demethylase 5A (KDM5A), KDM5B, KDM5C, KDM5D |
| Bosentan | Endothelin 1 |

**Supplementary Table 2 | Level 1 and level 2 drug dosing for male VICs cultured on stiff hydrogels.**

| Drug | EC1 (μM) | 5% Cmax (μM) | CC1 (μM) | Level 1 Dose | EC2 (μM) | 10% Cmax (μM) | CC2 (μM) | Level 2 Dose | Reference |
|---|---|---|---|---|---|---|---|---|---|
| Y-27632 | 0.558 | - | 10.8 | **0.558** | 1.34 | - | 19.9 | **1.34** | |
| LY294002 | 12.8 | - | 21.5 | **12.8** | 17.6 | - | 25.2 | **17.6** | |
| H1152 | 1.96 | - | 0.162 | **0.162** | 2.82 | - | 0.28 | **0.28** | |
| SB203580 | 24.8 | - | 0.413 | **0.413** | 40.8 | - | 2.18 | **2.18** | |
| Irosustat | 92.6 | 0.0106 | 48.5 | **0.0106** | 106.3 | 0.0212 | 58.7 | **0.0212** | [72] |
| TM5441 | 4.96 | - | 18.9 | **4.96** | 9.3 | - | 21.9 | **9.3** | |
| Losartan | 18.7 | 0.0265 | 378 | **0.0265** | 32.9 | 0.053 | 470 | **0.053** | [73] |
| SD-208 | 0.0582 | - | 26.6 | **0.0582** | 0.188 | - | 31.1 | **0.188** | |

**Supplementary Table 3 | Level 1 and level 2 drug dosing for female VICs cultured on stiff hydrogels.**

| Drug | EC1 (μM) | 5% Cmax (μM) | CC1 (μM) | Level 1 Dose | EC2 (μM) | 10% Cmax (μM) | CC2 (μM) | Level 2 Dose | Reference |
|---|---|---|---|---|---|---|---|---|---|
| Y-27632 | 1.31 | - | 27.5 | **1.31** | 2.81 | - | 37.9 | **2.81** | |
| LY294002 | 8.63 | - | 17.7 | **8.63** | 13.4 | - | 21.3 | **13.4** | |
| H1152 | 0.0765 | - | 0.26 | **0.0765** | 0.198 | - | 0.806 | **0.198** | |
| SB203580 | 7.56 | - | 29.8 | **7.56** | 14.5 | - | 48.9 | **14.5** | |
| Irosustat | 74.4 | 0.0106 | 50.2 | **0.0106** | 84.7 | 0.0212 | 57.9 | **0.0212** | [72] |
| TM5441 | 24.8 | - | 19.7 | **19.7** | 33.6 | - | 31.1 | **31.1** | |
| Losartan | 154 | 0.0265 | 11.1 | **0.0265** | 219 | 0.053 | 27 | **0.053** | [73] |
| SD-208 | 0.0061 | - | 34.8 | **0.0061** | 0.0275 | - | 40.1 | **0.0275** | |

**Supplementary Table 4 | 8-drug resolution IV orthogonal array composite design (OACD). The -1, 0, and 1 in the OACD represent the level 0, level 1, and level 2 concentrations, respectively.**

| Combination | Y-27632 | LY294002 | H1152 | SB203580 | Irosustat | TM5441 | Losartan | SD-208 |
|---|---|---|---|---|---|---|---|---|
| 1 | -1 | -1 | -1 | -1 | -1 | 1 | -1 | -1 |
| 2 | 1 | -1 | -1 | -1 | -1 | -1 | 1 | 1 |
| 3 | -1 | 1 | -1 | -1 | -1 | -1 | 1 | -1 |
| 4 | 1 | 1 | -1 | -1 | -1 | 1 | -1 | 1 |
| 5 | -1 | -1 | 1 | -1 | -1 | -1 | -1 | 1 |
| 6 | 1 | -1 | 1 | -1 | -1 | 1 | 1 | -1 |
| 7 | -1 | 1 | 1 | -1 | -1 | 1 | 1 | 1 |
| 8 | 1 | 1 | 1 | -1 | -1 | -1 | -1 | -1 |
| 9 | -1 | -1 | -1 | 1 | -1 | -1 | -1 | -1 |
| 10 | 1 | -1 | -1 | 1 | -1 | 1 | 1 | 1 |
| 11 | -1 | 1 | -1 | 1 | -1 | 1 | 1 | -1 |
| 12 | 1 | 1 | -1 | 1 | -1 | -1 | -1 | 1 |
| 13 | -1 | -1 | 1 | 1 | -1 | 1 | -1 | 1 |
| 14 | 1 | -1 | 1 | 1 | -1 | -1 | 1 | -1 |
| 15 | -1 | 1 | 1 | 1 | -1 | -1 | 1 | 1 |
| 16 | 1 | 1 | 1 | 1 | -1 | 1 | -1 | -1 |
| 17 | -1 | -1 | -1 | -1 | 1 | 1 | 1 | 1 |
| 18 | 1 | -1 | -1 | -1 | 1 | -1 | -1 | -1 |
| 19 | -1 | 1 | -1 | -1 | 1 | -1 | -1 | 1 |
| 20 | 1 | 1 | -1 | -1 | 1 | 1 | 1 | -1 |
| 21 | -1 | -1 | 1 | -1 | 1 | -1 | 1 | -1 |
| 22 | 1 | -1 | 1 | -1 | 1 | 1 | -1 | 1 |
| 23 | -1 | 1 | 1 | -1 | 1 | 1 | -1 | -1 |
| 24 | 1 | 1 | 1 | -1 | 1 | -1 | 1 | 1 |
| 25 | -1 | -1 | -1 | 1 | 1 | -1 | 1 | 1 |
| 26 | 1 | -1 | -1 | 1 | 1 | 1 | -1 | -1 |
| 27 | -1 | 1 | -1 | 1 | 1 | 1 | -1 | 1 |
| 28 | 1 | 1 | -1 | 1 | 1 | -1 | 1 | -1 |
| 29 | -1 | -1 | 1 | 1 | 1 | 1 | 1 | -1 |
| 30 | 1 | -1 | 1 | 1 | 1 | -1 | -1 | 1 |
| 31 | -1 | 1 | 1 | 1 | 1 | -1 | -1 | -1 |
| 32 | 1 | 1 | 1 | 1 | 1 | 1 | 1 | 1 |
| 33 | -1 | -1 | -1 | -1 | -1 | -1 | -1 | -1 |
| 34 | -1 | 0 | 0 | -1 | -1 | 0 | 1 | 1 |
| 35 | -1 | 1 | 1 | -1 | -1 | 1 | 0 | 0 |
| 36 | -1 | -1 | 0 | 1 | 0 | -1 | 0 | 0 |
| 37 | -1 | 0 | 1 | 1 | 0 | 0 | -1 | -1 |
| 38 | -1 | 1 | -1 | 1 | 0 | 1 | 1 | 1 |
| 39 | -1 | -1 | 1 | 0 | 1 | -1 | 1 | 1 |
| 40 | -1 | 0 | -1 | 0 | 1 | 0 | 0 | 0 |
| 41 | -1 | 1 | 0 | 0 | 1 | 1 | -1 | -1 |
| 42 | 0 | -1 | 0 | 0 | -1 | 0 | -1 | 0 |
| 43 | 0 | 0 | 1 | 0 | -1 | 1 | 1 | -1 |
| 44 | 0 | 1 | -1 | 0 | -1 | -1 | 0 | 1 |
| 45 | 0 | -1 | 1 | -1 | 0 | 0 | 0 | 1 |
| 46 | 0 | 0 | -1 | -1 | 0 | 1 | -1 | 0 |
| 47 | 0 | 1 | 0 | -1 | 0 | -1 | 1 | -1 |
| 48 | 0 | -1 | -1 | 1 | 1 | 0 | 1 | -1 |
| 49 | 0 | 0 | 0 | 1 | 1 | 1 | 0 | 1 |
| 50 | 0 | 1 | 1 | 1 | 1 | -1 | -1 | 0 |
| 51 | 1 | -1 | 1 | 1 | -1 | 1 | -1 | 1 |
| 52 | 1 | 0 | -1 | 1 | -1 | -1 | 1 | 0 |
| 53 | 1 | 1 | 0 | 1 | -1 | 0 | 0 | -1 |
| 54 | 1 | -1 | -1 | 0 | 0 | 1 | 0 | -1 |
| 55 | 1 | 0 | 0 | 0 | 0 | -1 | -1 | 1 |
| 56 | 1 | 1 | 1 | 0 | 0 | 0 | 1 | 0 |
| 57 | 1 | -1 | 0 | -1 | 1 | 1 | 1 | 0 |
| 58 | 1 | 0 | 1 | -1 | 1 | -1 | 0 | -1 |
| 59 | 1 | 1 | -1 | -1 | 1 | 0 | -1 | 1 |

**Supplementary Table 5 | Percent αSMA reduction for all 59 combinations used to generate the IDentif.AI model fit for male VICs cultured on hydrogels. n= 3-4 gels.**

| Combination | Replicate 1 | Replicate 2 | Replicate 3 | Replicate 4 | Average | Standard Deviation | Standard Error |
|---|---|---|---|---|---|---|---|
| 1 | -10.73 | 16.81 | 36.20 | 30.17 | 18.11 | 20.86 | 10.43 |
| 2 | 46.82 | 27.64 | 33.34 | 16.42 | 31.06 | 12.64 | 6.32 |
| 3 | 31.59 | 42.82 | 34.98 | 53.51 | 40.72 | 9.74 | 4.87 |
| 4 | 64.89 | 36.33 | 28.66 | - | 43.29 | 19.09 | 11.02 |
| 5 | -1.51 | 15.24 | -8.39 | 2.32 | 1.91 | 9.93 | 4.96 |
| 6 | 0.97 | 17.38 | 17.78 | - | 12.04 | 9.59 | 5.54 |
| 7 | 59.68 | 57.16 | 32.89 | 37.34 | 46.77 | 13.61 | 6.81 |
| 8 | 6.05 | 14.17 | 12.02 | 2.83 | 8.77 | 5.24 | 2.62 |
| 9 | 2.77 | 17.21 | 28.83 | 29.22 | 19.51 | 12.47 | 6.24 |
| 10 | 47.60 | 53.29 | 48.97 | 36.33 | 46.55 | 7.23 | 3.62 |
| 11 | -1.57 | -2.70 | -0.50 | - | -1.59 | 1.10 | 0.63 |
| 12 | -12.34 | 27.59 | 39.04 | 28.04 | 20.58 | 22.58 | 11.29 |
| 13 | 39.66 | 16.08 | 17.49 | 47.14 | 30.09 | 15.67 | 7.84 |
| 14 | 13.21 | 19.58 | 31.93 | 9.09 | 18.45 | 9.97 | 4.98 |
| 15 | -8.79 | 10.50 | 5.82 | 19.58 | 6.78 | 11.85 | 5.92 |
| 16 | 44.99 | 40.00 | 36.44 | 31.82 | 38.31 | 5.57 | 2.79 |
| 17 | 7.74 | -6.64 | -6.87 | 26.52 | 5.19 | 15.78 | 7.89 |
| 18 | 11.12 | 11.01 | 36.56 | 18.90 | 19.40 | 12.02 | 6.01 |
| 19 | 19.69 | -5.01 | 10.56 | -2.30 | 5.74 | 11.52 | 5.76 |
| 20 | 45.60 | 27.76 | 35.99 | 14.34 | 30.92 | 13.25 | 6.62 |
| 21 | 0.58 | 19.47 | 30.75 | - | 16.93 | 15.24 | 8.80 |
| 22 | 39.66 | 33.12 | 21.27 | 40.45 | 33.62 | 8.87 | 4.43 |
| 23 | 58.14 | 50.08 | 50.86 | - | 53.03 | 4.44 | 2.56 |
| 24 | 51.11 | 36.03 | 39.64 | - | 42.26 | 7.88 | 4.55 |
| 25 | 19.76 | 18.42 | 27.46 | 31.54 | 24.29 | 6.26 | 3.13 |
| 26 | 54.85 | 37.84 | 36.44 | 3.08 | 33.05 | 21.66 | 10.83 |
| 27 | -1.58 | 19.53 | -24.74 | -2.81 | -2.40 | 18.08 | 9.04 |
| 28 | 9.15 | 15.85 | 24.66 | 46.25 | 23.98 | 16.15 | 8.08 |
| 29 | 55.38 | 39.88 | 12.99 | 6.70 | 28.74 | 22.86 | 11.43 |
| 30 | 19.00 | 10.90 | 20.11 | - | 16.67 | 5.03 | 2.90 |
| 31 | -1.88 | 13.17 | 10.37 | - | 7.22 | 8.00 | 4.62 |
| 32 | 53.52 | 50.14 | 41.22 | 35.50 | 45.10 | 8.24 | 4.12 |
| 33 | 24.72 | 14.98 | 5.30 | -26.60 | 4.60 | 22.26 | 11.13 |
| 34 | 3.84 | 24.95 | 23.26 | - | 17.35 | 11.73 | 6.77 |
| 35 | 50.48 | 35.56 | 37.14 | - | 41.06 | 8.20 | 4.73 |
| 36 | 7.28 | 14.22 | 6.29 | 16.26 | 11.01 | 4.97 | 2.48 |
| 37 | -0.42 | 29.96 | 31.95 | 29.32 | 22.70 | 15.45 | 7.73 |
| 38 | 50.77 | 32.53 | 54.93 | 4.48 | 35.68 | 22.96 | 11.48 |
| 39 | 7.51 | 4.54 | -11.03 | - | 0.34 | 9.96 | 5.75 |
| 40 | -4.50 | 35.68 | -3.92 | - | 9.09 | 23.03 | 13.30 |
| 41 | 33.87 | 27.81 | 34.51 | 23.61 | 29.95 | 5.20 | 2.60 |
| 42 | 29.61 | 25.24 | 19.88 | 16.90 | 22.91 | 5.65 | 2.82 |
| 43 | 22.21 | 13.64 | -4.62 | 3.20 | 8.61 | 11.75 | 5.88 |
| 44 | 42.68 | 42.13 | 48.86 | 32.76 | 41.61 | 6.64 | 3.32 |
| 45 | 55.72 | 53.68 | 47.57 | - | 52.32 | 4.24 | 2.45 |
| 46 | 58.73 | 37.46 | 25.79 | 33.94 | 38.98 | 14.04 | 7.02 |
| 47 | 36.00 | 30.84 | 42.89 | 58.53 | 42.06 | 12.04 | 6.02 |
| 48 | 20.30 | 21.25 | 5.09 | -7.08 | 9.89 | 13.52 | 6.76 |
| 49 | 43.55 | 39.34 | 37.05 | 43.37 | 40.83 | 3.18 | 1.59 |
| 50 | 14.53 | 24.68 | 24.62 | 30.42 | 23.56 | 6.61 | 3.31 |
| 51 | 31.66 | 20.74 | 15.72 | 16.61 | 21.18 | 7.32 | 3.66 |
| 52 | 35.00 | 39.87 | 24.91 | 18.11 | 29.47 | 9.81 | 4.90 |
| 53 | 13.59 | 25.14 | 34.65 | 42.27 | 28.91 | 12.39 | 6.19 |
| 54 | 20.89 | 23.25 | 9.82 | -11.44 | 10.63 | 15.83 | 7.92 |
| 55 | 26.85 | -4.60 | 28.61 | -7.82 | 10.76 | 19.65 | 9.83 |
| 56 | 6.20 | 6.90 | 0.92 | - | 4.67 | 3.27 | 1.89 |
| 57 | 17.20 | 15.57 | -1.33 | -3.10 | 7.08 | 10.78 | 5.39 |
| 58 | 28.84 | 11.18 | -0.67 | 8.07 | 11.86 | 12.38 | 6.19 |
| 59 | 23.15 | -3.37 | -19.73 | 3.32 | 0.84 | 17.75 | 8.87 |

**Supplementary Table 6 | Percent αSMA reduction for individual level 1 and level 2 drug doses used to generate the IDentif.AI model fit for male VICs cultured on stiff hydrogels. n= 3-4 gels. SD=Standard deviation. SE=Standard error.**

| | Drug | Replicate 1 | Replicate 2 | Replicate 3 | Replicate 4 | Average | SD | SE |
|---|---|---|---|---|---|---|---|---|
| Level 1 dose | Y-27632 | 12.82 | -12.17 | 2.34 | -2.04 | 0.24 | 10.36 | 5.18 |
| | LY294002 | -9.56 | 18.87 | 11.27 | - | 6.86 | 14.72 | 8.50 |
| | H1152 | -2.68 | -10.05 | -14.68 | 7.31 | -5.03 | 9.59 | 4.80 |
| | SB203580 | 18.18 | 8.93 | -10.77 | - | 5.45 | 14.78 | 8.53 |
| | Irosustat | 6.97 | 1.96 | -6.40 | - | 0.84 | 6.75 | 3.90 |
| | TM5441 | -3.67 | 25.74 | -26.46 | - | -1.46 | 26.17 | 15.11 |
| | Losartan | 1.05 | 10.96 | 7.66 | 2.66 | 5.58 | 4.56 | 2.28 |
| | SD-208 | 31.08 | -7.59 | 20.32 | - | 14.60 | 19.96 | 11.53 |
| Level 2 dose | Y-27632 | 30.83 | -6.93 | -11.04 | -3.38 | 2.37 | 19.23 | 9.62 |
| | LY294002 | 7.61 | 0.98 | -3.19 | 16.60 | 5.50 | 8.63 | 4.32 |
| | H1152 | 23.58 | 3.59 | -18.16 | -6.37 | 0.66 | 17.68 | 8.84 |
| | SB203580 | 9.92 | -17.32 | 26.88 | -4.78 | 3.67 | 19.06 | 9.53 |
| | Irosustat | 0.11 | -17.93 | 13.54 | 6.97 | 0.67 | 13.56 | 6.78 |
| | TM5441 | 13.93 | 0.51 | 6.21 | -11.91 | 2.18 | 10.89 | 5.44 |
| | Losartan | -2.70 | 1.72 | 5.92 | -0.97 | 0.99 | 3.76 | 1.88 |
| | SD-208 | 14.55 | -15.09 | -11.20 | 14.27 | 0.63 | 15.99 | 7.99 |

**Supplementary Table 7 | Coefficients and model fit parameters used for IDentif.AI analysis for male VICs cultured on stiff hydrogels. All terms are shown after a square transformation to improve the overall model fit. Statistical significance determined by F-test. *=P<0.05, **=P<0.01, ***=P<0.001, ****=P<0.0001.**

| Term | Estimate | Statistical Significance |
|---|---|---|
| Intercept | 1380.2 | **** |
| Y-27632 | 177.10 | * |
| LY294002 | 250.12 | *** |
| H1152 | 132.26 | ns |
| SB203580 | -82.663 | ns |
| Irosustat | 26.218 | ns |
| TM5441 | 355.91 | **** |
| Losartan | 77.384 | ns |
| SD-208 | 183.28 | * |
| Y-27632:SB203580 | 172.55 | * |
| LY294002:SB203580 | -258.25 | ** |
| LY294002:TM5441 | 140.01 | ns |
| H1152:SB203580 | -156.00 | * |
| H1152:Irosustat | 276.25 | *** |
| H1152:TM5441 | 225.02 | ** |
| Irosustat:SD-208 | -179.49 | * |
| TM5441:Losartan | -212.45 | ** |
| Losartan:SD-208 | 253.83 | ** |
| Y-27632$^2$ | -766.35 | *** |
| SB203580$^2$ | 634.05 | ** |
| Irosustat$^2$ | -349.22 | ns |
| Degrees of Freedom | 54 | |
| $R^2$ | 0.748 | |
| Adjusted $R^2$ | 0.655 | |
| F-statistic vs constant model | 8.02 | **** |

**Supplementary Table 8 | Top IDentif.AI predicted effective 2-drug, 3-drug, and 4-drug combinations and two low ranked drug combinations for male VICs selected for *in vitro* validation. Drug doses are indicated as -1 for no drug, 0 for level 1 dose, and 1 for level 2 dose. Abbreviations: Y-27632 (Y27); LY294002 (LY); H1152 (H11); SB203580 (SB); Irosustat (Iro); TM5441 (TM); Losartan (Los); SD-208 (SD).**

| Combo # | Y27 | LY | H11 | SB | Iro | TM | Los | SD | Predicted % αSMA Reduction |
|---|---|---|---|---|---|---|---|---|---|
| M1 | -1 | 1 | -1 | -1 | -1 | 1 | -1 | -1 | 36.72 |
| M2 | 0 | 1 | -1 | -1 | -1 | -1 | -1 | -1 | 34.31 |
| M3 | -1 | -1 | -1 | -1 | -1 | -1 | 1 | 1 | 31.65 |
| M4 | 0 | 1 | -1 | -1 | -1 | 1 | -1 | -1 | 45.11 |
| M5 | -1 | 1 | 1 | -1 | -1 | 1 | -1 | -1 | 42.06 |
| M6 | -1 | 1 | -1 | -1 | -1 | -1 | 1 | 1 | 40.60 |
| M7 | -1 | 1 | 1 | -1 | 1 | 1 | -1 | -1 | 51.37 |
| M8 | 0 | 1 | 1 | -1 | -1 | 1 | -1 | -1 | 49.64 |
| M9 | 0 | 1 | -1 | -1 | -1 | -1 | 1 | 1 | 48.38 |
| M10 | -1 | 1 | -1 | 1 | -1 | -1 | -1 | -1 | 2.86 |
| M11 | 1 | -1 | 1 | 1 | -1 | -1 | -1 | -1 | -2.23 |

**Supplementary Table 9 | Percent αSMA reduction for all 59 combinations used to generate the IDentif.AI model fit for female VICs cultured on hydrogels. n = 2-4 gels.**

| Combination | Replicate 1 | Replicate 2 | Replicate 3 | Replicate 4 | Average | Standard Deviation | Standard Error |
|---|---|---|---|---|---|---|---|
| 1 | 29.24 | 24.65 | 31.88 | - | 28.59 | 3.65 | 2.11 |
| 2 | 34.90 | 41.86 | 43.89 | - | 40.22 | 4.71 | 2.72 |
| 3 | -5.17 | 21.00 | 22.53 | 38.61 | 19.24 | 18.12 | 9.06 |
| 4 | 51.11 | 49.67 | 41.40 | 40.55 | 45.68 | 5.48 | 2.74 |
| 5 | 24.36 | 35.36 | 28.29 | 20.31 | 27.08 | 6.41 | 3.20 |
| 6 | 36.61 | 47.31 | 44.28 | 44.87 | 43.27 | 4.63 | 2.31 |
| 7 | -5.17 | 19.86 | 24.81 | 27.72 | 16.81 | 15.01 | 7.50 |
| 8 | 30.46 | 33.54 | 29.26 | 9.42 | 25.67 | 10.98 | 5.49 |
| 9 | 2.19 | -7.11 | 24.36 | 24.81 | 11.06 | 16.07 | 8.04 |
| 10 | 44.60 | 41.80 | 35.87 | 15.64 | 34.48 | 13.08 | 6.54 |
| 11 | -2.09 | 6.23 | 19.68 | 22.31 | 11.53 | 11.49 | 5.75 |
| 12 | 36.04 | 39.58 | 36.22 | 36.96 | 37.20 | 1.64 | 0.82 |
| 13 | 14.50 | -15.20 | 21.79 | - | 7.03 | 19.59 | 11.31 |
| 14 | -7.11 | 14.78 | -0.44 | 5.15 | 3.10 | 9.26 | 4.63 |
| 15 | 37.24 | 23.67 | -1.18 | 20.25 | 20.00 | 15.91 | 7.95 |
| 16 | 43.71 | 56.04 | 51.32 | - | 50.36 | 6.22 | 3.59 |
| 17 | 7.09 | -18.11 | -11.61 | - | -7.54 | 13.08 | 7.55 |
| 18 | 18.09 | 34.11 | 51.14 | 22.53 | 31.47 | 14.75 | 7.37 |
| 19 | 11.99 | 15.81 | -4.66 | - | 7.71 | 10.88 | 6.28 |
| 20 | 25.61 | -11.72 | -13.32 | - | 0.19 | 22.03 | 12.72 |
| 21 | 28.80 | 19.97 | 1.16 | 14.78 | 16.18 | 11.57 | 5.78 |
| 22 | 18.43 | 27.89 | 6.00 | - | 17.44 | 10.98 | 6.34 |
| 23 | 41.47 | 57.08 | 66.18 | - | 54.91 | 12.50 | 7.22 |
| 24 | 42.63 | 37.20 | 29.71 | - | 36.51 | 6.48 | 3.74 |
| 25 | 32.76 | 14.86 | 32.94 | - | 26.86 | 10.39 | 6.00 |
| 26 | 43.91 | 47.85 | 36.36 | - | 42.71 | 5.84 | 3.37 |
| 27 | 28.57 | 15.88 | 22.83 | - | 22.43 | 6.36 | 3.67 |
| 28 | 46.38 | 39.47 | 44.28 | - | 43.38 | 3.54 | 2.05 |
| 29 | 47.79 | 45.74 | 46.70 | 23.84 | 41.02 | 11.48 | 5.74 |
| 30 | 41.70 | 30.31 | 11.69 | 29.71 | 28.35 | 12.40 | 6.20 |
| 31 | 28.69 | 18.16 | 34.02 | 8.46 | 22.33 | 11.36 | 5.68 |
| 32 | 42.41 | 48.94 | 40.33 | - | 43.89 | 4.49 | 2.59 |
| 33 | 2.65 | -5.55 | 5.28 | -3.22 | -0.21 | 5.03 | 2.52 |
| 34 | 20.31 | 6.72 | 19.77 | 13.49 | 15.07 | 6.37 | 3.19 |
| 35 | 35.52 | 15.58 | 9.95 | 37.97 | 24.76 | 14.07 | 7.03 |
| 36 | 3.25 | 5.70 | 0.61 | -6.63 | 0.73 | 5.33 | 2.67 |
| 37 | 23.66 | 25.04 | 26.78 | 10.13 | 21.40 | 7.62 | 3.81 |
| 38 | 15.10 | 29.47 | -5.19 | - | 13.13 | 17.42 | 10.06 |
| 39 | -2.08 | 9.59 | 5.52 | - | 4.35 | 5.93 | 3.42 |
| 40 | -12.32 | -8.33 | 18.54 | - | -0.70 | 16.79 | 9.69 |
| 41 | 16.24 | 29.41 | 11.33 | - | 18.99 | 9.35 | 5.40 |
| 42 | 24.38 | 27.80 | 28.33 | 23.78 | 26.07 | 2.32 | 1.16 |
| 43 | 26.24 | 19.11 | 12.17 | 25.16 | 20.67 | 6.48 | 3.24 |
| 44 | 30.13 | 45.02 | 30.91 | - | 35.35 | 8.38 | 4.84 |
| 45 | 15.40 | 7.37 | 31.28 | - | 18.02 | 12.17 | 7.02 |
| 46 | 29.92 | 21.96 | 45.56 | - | 32.48 | 12.00 | 6.93 |
| 47 | 34.06 | 41.16 | 45.10 | - | 40.11 | 5.60 | 3.23 |
| 48 | 28.20 | 22.25 | 18.10 | 17.15 | 21.42 | 5.03 | 2.52 |
| 49 | 36.18 | 30.52 | 31.38 | 21.01 | 29.77 | 6.35 | 3.17 |
| 50 | 38.03 | 38.05 | 46.46 | - | 40.85 | 4.86 | 2.81 |
| 51 | -8.45 | 5.04 | 9.64 | 19.19 | 6.35 | 11.50 | 5.75 |
| 52 | 37.54 | 17.88 | 23.86 | 19.41 | 24.67 | 8.95 | 4.47 |
| 53 | 4.24 | 40.01 | - | - | 22.13 | 25.30 | 17.89 |
| 54 | 27.45 | 29.07 | 18.39 | - | 24.97 | 5.76 | 3.32 |
| 55 | 11.17 | 14.81 | 26.05 | - | 17.34 | 7.75 | 4.48 |
| 56 | 36.58 | 55.48 | - | - | 46.03 | 13.37 | 9.45 |
| 57 | -17.06 | 0.08 | -1.60 | - | -6.19 | 9.45 | 5.46 |
| 58 | 23.42 | 20.58 | 40.44 | - | 28.14 | 10.74 | 6.20 |
| 59 | 37.53 | 21.09 | - | - | 29.31 | 11.63 | 8.22 |

**Supplementary Table 10 | Percent αSMA reduction for individual level 1 and level 2 drug doses used to generate the IDentif.AI model fit for female VICs cultured on stiff hydrogels. n= 3-4 gels. SD=Standard deviation. SE=Standard error.**

| | Drug | Replicate 1 | Replicate 2 | Replicate 3 | Replicate 4 | Average | SD | SE |
|---|---|---|---|---|---|---|---|---|
| Level 1 dose | Y-27632 | 26.45 | 26.29 | 34.04 | | 28.93 | 4.43 | 2.56 |
| | LY294002 | 17.97 | 11.18 | 18.50 | -9.65 | 9.50 | 13.19 | 6.60 |
| | H1152 | 11.18 | 6.02 | 6.93 | - | 8.04 | 2.76 | 1.59 |
| | SB203580 | 9.86 | 4.49 | -7.28 | - | 2.35 | 8.77 | 5.06 |
| | Irosustat | 27.17 | 7.47 | -5.77 | - | 9.62 | 16.58 | 9.57 |
| | TM5441 | 2.03 | -0.44 | 17.70 | -13.20 | 1.52 | 12.68 | 6.34 |
| | Losartan | 24.32 | -7.12 | 3.81 | - | 7.00 | 15.96 | 9.21 |
| | SD-208 | -7.55 | 18.40 | 1.71 | - | 4.19 | 13.15 | 7.59 |
| Level 2 dose | Y-27632 | 22.31 | 22.17 | 21.88 | - | 22.12 | 0.22 | 0.13 |
| | LY294002 | -9.86 | 20.44 | -20.52 | 20.93 | 2.75 | 21.17 | 10.58 |
| | H1152 | 24.96 | 11.99 | 2.25 | 10.38 | 12.39 | 9.40 | 4.70 |
| | SB203580 | 5.32 | -22.94 | -0.01 | -8.57 | -6.55 | 12.33 | 6.17 |
| | Irosustat | -2.33 | -6.74 | 7.41 | - | -0.55 | 7.24 | 4.18 |
| | TM5441 | -4.05 | -2.76 | -16.11 | - | -7.64 | 7.36 | 4.25 |
| | Losartan | 6.05 | -1.18 | 7.70 | -3.07 | 2.37 | 5.30 | 2.65 |
| | SD-208 | 8.87 | 12.26 | 5.32 | - | 8.81 | 3.47 | 2.00 |

**Supplementary Table 11 | Coefficients and model fit parameters used for IDentif.AI analysis for female VICs cultured on stiff hydrogels. Statistical significance determined by F-test. *=P<0.05, **=P<0.01, ***=P<0.001, ****=P<0.0001.**

| Term | Estimate | Statistical Significance |
|---|---|---|
| Intercept | 23.784 | **** |
| Y-27632 | 7.493 | **** |
| LY294002 | 4.8144 | *** |
| H1152 | 2.6284 | * |
| SB203580 | 1.4806 | ns |
| Irosustat | 1.4956 | ns |
| TM5441 | 1.1921 | ns |
| Losartan | -1.636 | ns |
| SD-208 | -0.50618 | ns |
| Y-27632:H1152 | -3.944 | ** |
| LY294002:H1152 | 2.7807 | * |
| H1152:SB203580 | -3.0194 | * |
| H1152:Irosustat | 2.8757 | * |
| H1152:TM5441 | 2.7529 | * |
| H1152:SD-208 | -3.6649 | ** |
| SB203580:Irosustat | 5.0124 | *** |
| SB203580:TM5441 | 2.7493 | * |
| TM5441:Losartan | -3.5073 | ** |
| TM5441:SD-208 | -2.7553 | * |
| $Y\text{-}27632^2$ | -7.5615 | * |
| $H\text{-}1152^2$ | 6.7199 | * |
| Degrees of Freedom | 54 | |
| $R^2$ | 0.745 | |
| Adjusted $R^2$ | 0.651 | |
| F-statistic vs constant model | 7.89 | **** |

**Supplementary Table 12 | Top IDentif.AI predicted effective 2-drug, 3-drug, and 4-drug combinations and three low ranked drug combinations for female VICs selected for *in vitro* validation. Drug doses are indicated as -1 for no drug, 0 for level 1 dose, and 1 for level 2 dose. Abbreviations: Y-27632 (Y27); LY294002 (LY); H1152 (H11); SB203580 (SB); Irosustat (Iro); TM5441 (TM); Losartan (Los); SD-208 (SD).**

| Combo # | Y27 | LY | H11 | SB | Iro | TM | Los | SD | Predicted % αSMA Reduction |
|---|---|---|---|---|---|---|---|---|---|
| F1 | 1 | -1 | -1 | -1 | -1 | -1 | -1 | 1 | 39.96 |
| F2 | -1 | 1 | 1 | -1 | -1 | -1 | -1 | -1 | 30.15 |
| F3 | -1 | -1 | 1 | -1 | -1 | 1 | -1 | -1 | 29.87 |
| F4 | -1 | 1 | 1 | -1 | -1 | 1 | -1 | -1 | 45.06 |
| F5 | 1 | 1 | -1 | -1 | -1 | -1 | -1 | 1 | 44.03 |
| F6 | 1 | -1 | -1 | -1 | -1 | -1 | 1 | 1 | 43.70 |
| F7 | 0 | 1 | 1 | -1 | -1 | 1 | -1 | -1 | 56.17 |
| F8 | 1 | 1 | -1 | -1 | -1 | -1 | 1 | 1 | 47.77 |
| F9 | -1 | 1 | 1 | -1 | 0 | 1 | -1 | -1 | 44.42 |
| F10 | -1 | -1 | -1 | -1 | 1 | 0 | -1 | -1 | -5.57 |
| F11 | -1 | -1 | 1 | 1 | -1 | -1 | -1 | 1 | -6.48 |
| F12 | -1 | -1 | -1 | -1 | 1 | 1 | 1 | 0 | -13.50 |

**Supplementary Table 13 | Top IDentif.AI predicted differentially effective 2-drug, 3-drug, and 4-drug combinations. Drug doses are indicated as -1 for no drug, 0 for level 1 dose, and 1 for level 2 dose. Abbreviations: Y-27632 (Y27); LY294002 (LY); H1152 (H11); SB203580 (SB); Irosustat (Iro); TM5441 (TM); Losartan (Los); SD-208 (SD).**

| Combo # | Y27 | LY | H11 | SB | Iro | TM | Los | SD | Predicted Male-Female % αSMA Reduction |
|---|---|---|---|---|---|---|---|---|---|
| MB1 | -1 | **1** | -1 | -1 | **0** | -1 | -1 | -1 | 24.52 |
| MB2 | -1 | **1** | -1 | -1 | **1** | **1** | -1 | -1 | 34.55 |
| MB3 | -1 | **1** | -1 | -1 | -1 | **1** | **1** | **1** | 38.11 |
| FB1 | **1** | -1 | -1 | -1 | -1 | -1 | -1 | **1** | -28.64 |
| FB2 | **1** | -1 | -1 | **0** | -1 | -1 | -1 | **0** | -30.47 |
| FB3 | **1** | -1 | -1 | **0** | **0** | -1 | -1 | **1** | -32.79 |

**Supplementary Table 14 | Forward and reverse primers for RT-qPCR. All primers are shown from 5’-3’ end.**

| Gene | Forward Primer (5’-3’) | Reverse Primer (5’-3’) |
|---|---|---|
| *RPL30* | AGATTTCCTCAAGGCTGGGC | GCTGGGGTACAAGCAGACTC |
| *ACTA2* | GCAAACAGGAATACGATGAAGCC | AACACATAGGTAACGAGTCAGAGC |
| *COL1A1* | GGGCAAGACAGTGATTGAATACA | GGATGGAGGGAGTTTACAGGAA |

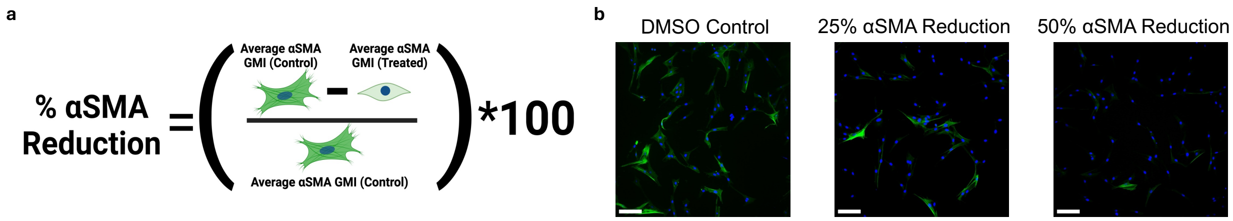


**Supplementary Fig. 1 | Overview of percent αSMA reduction output. a,** Schematic defining how percent αSMA reduction is calculated. GMI=Gradient mean intensity. **b,** Representative immunofluorescent images of male VICs cultured on hydrogels with αSMA stained in green and DAPI stained in blue. Scale bar = 100 μm.

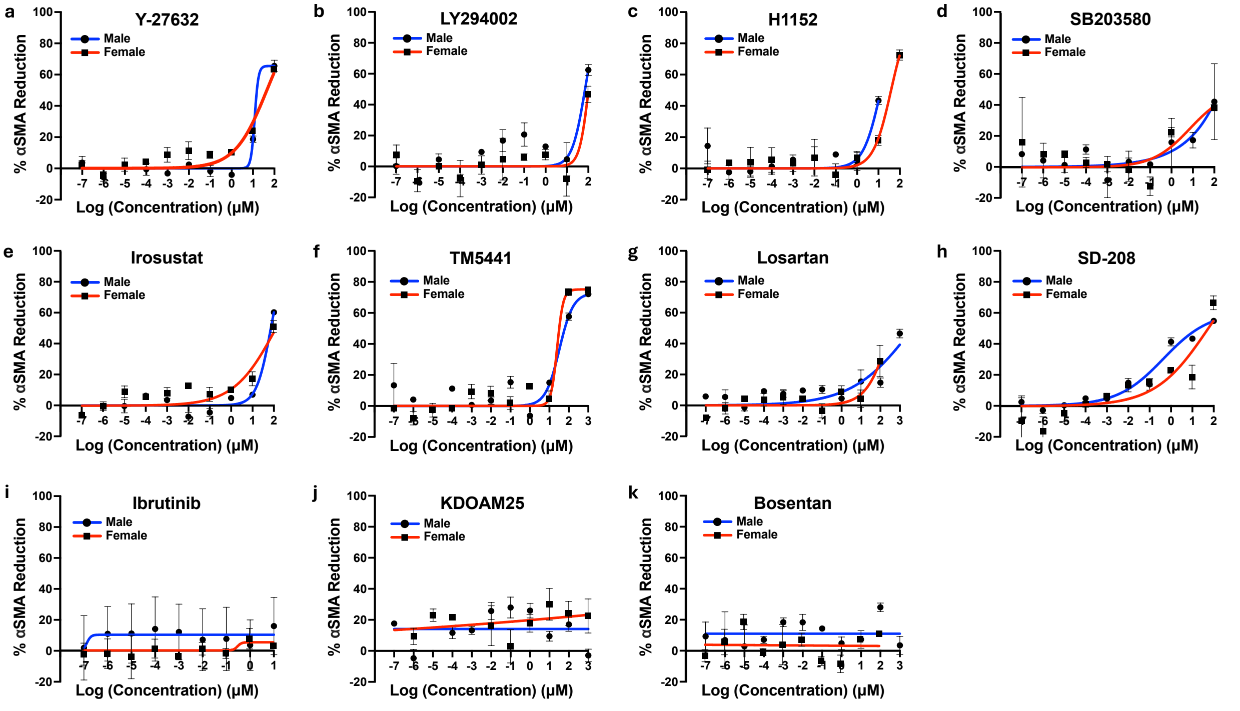


**Supplementary Fig. 2 | Eight out of eleven anti-fibrotic drugs inhibit myofibroblast activation in male and female VICs on TCPS. a-k,** Percent αSMA reduction in male and female VICs cultured on TCPS at doses ranging from $10^{-7}$ μM up to $10^{3}$ μM for **a** Y-27632, **b** LY294002, **c** H1152, **d** SB203580, **e** Irosustat, **f** TM5441, **g** Losartan, **h** SD-208, **i** Ibrutinib, **j** KDOAM25, and **k** Bosentan (n = 2 wells). Each data point is plotted as mean ± standard error of the mean. The best fit line shown was generated using a nonlinear regression curve fit with GraphPad Prism.

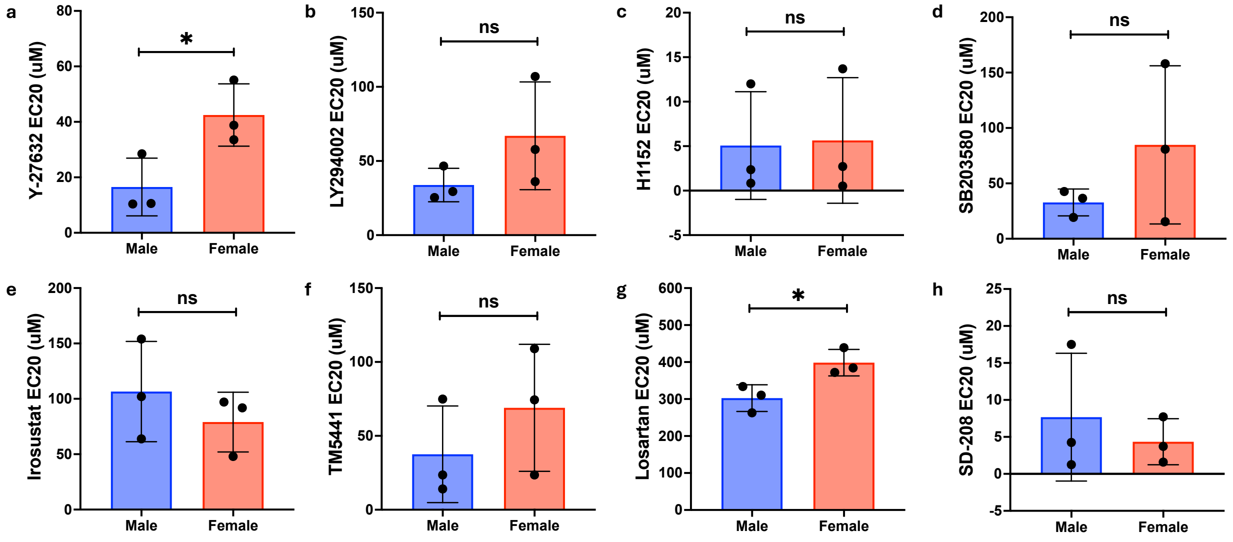


**Supplementary Fig. 3 | Male VICs cultured on TCPS are more responsive to Y-27632 and Losartan relative to female VICs. a-h,** Male versus female $EC_{20}$ values for VICs cultured on stiff hydrogels with **a** Y-27632, **b** LY294002, **c** H1152, **d** SB203580, **e** Irosustat, **f** TM5441, **g** Losartan, and **h** SD-208 (N = 3 biological replicates). Data is plotted as mean ± standard deviation. Statistical significance was determined by an unpaired two-tailed t-test with Welch's correction and indicated as *=P<0.05.

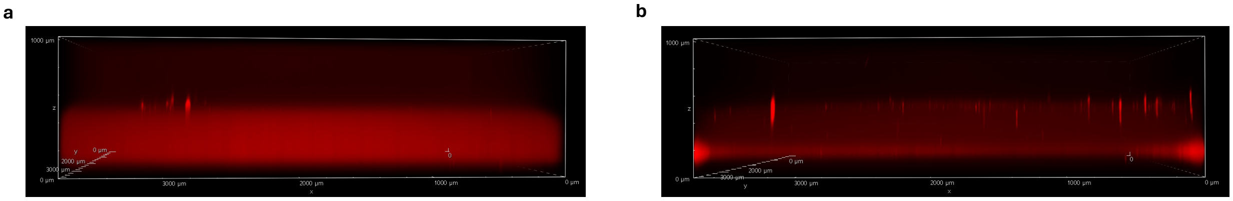


**Supplementary Fig. 4 | Soft and stiff hydrogels formed in a 96-well plate are flat.** **a,b,** Representative immunofluorescent side-view images of **a** soft and **b** stiff hydrogels stained with Cy5.

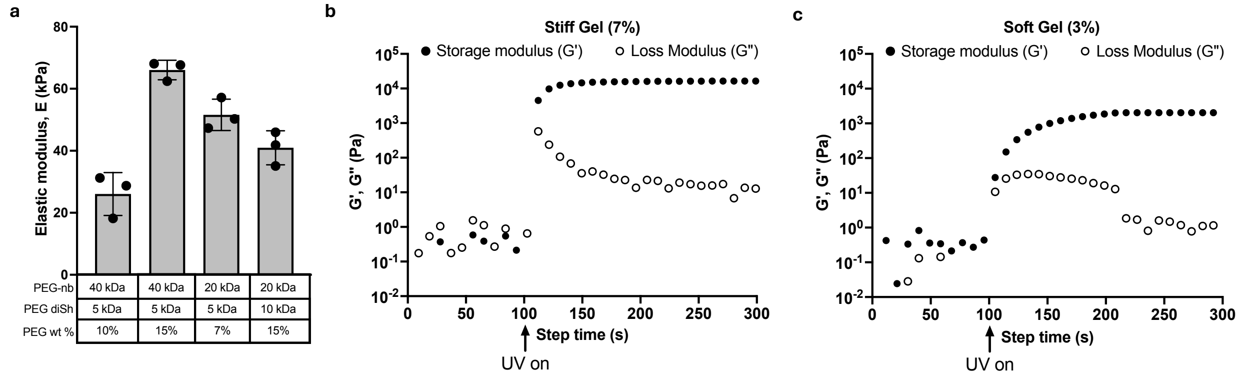


**Supplementary Fig. 5 | Rheological measurements of various hydrogel compositions. a,** Rheology of candidate stiff gels comprised of different PEG monomers. Data shown as mean ± standard deviation (n = 3 gel samples). **b,c,** Storage and loss moduli of optimized **b** stiff and **c** soft hydrogel precursor solution (n = 1; representative replicate shown).

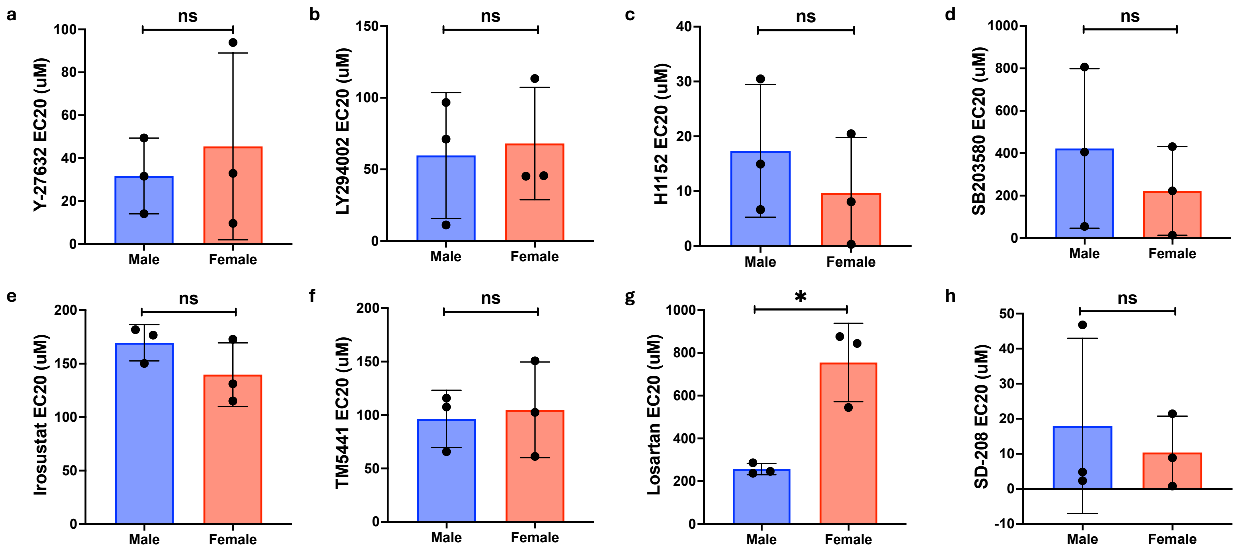


**Supplementary Fig. 6 | Male VICs cultured on stiff hydrogels are more responsive to Losartan relative to female VICs. a-h,** Male versus female $EC_{20}$ values for VICs cultured on stiff hydrogels with **a** Y-27632, **b** LY294002, **c** H1152, **d** SB203580, **e** Irosustat, **f** TM5441, **g** Losartan, and **h** SD-208 (N = 3 biological replicates). Data is plotted as mean ± standard deviation. Statistical significance was determined by an unpaired two-tailed t-test with Welch's correction and indicated as *=P<0.05.

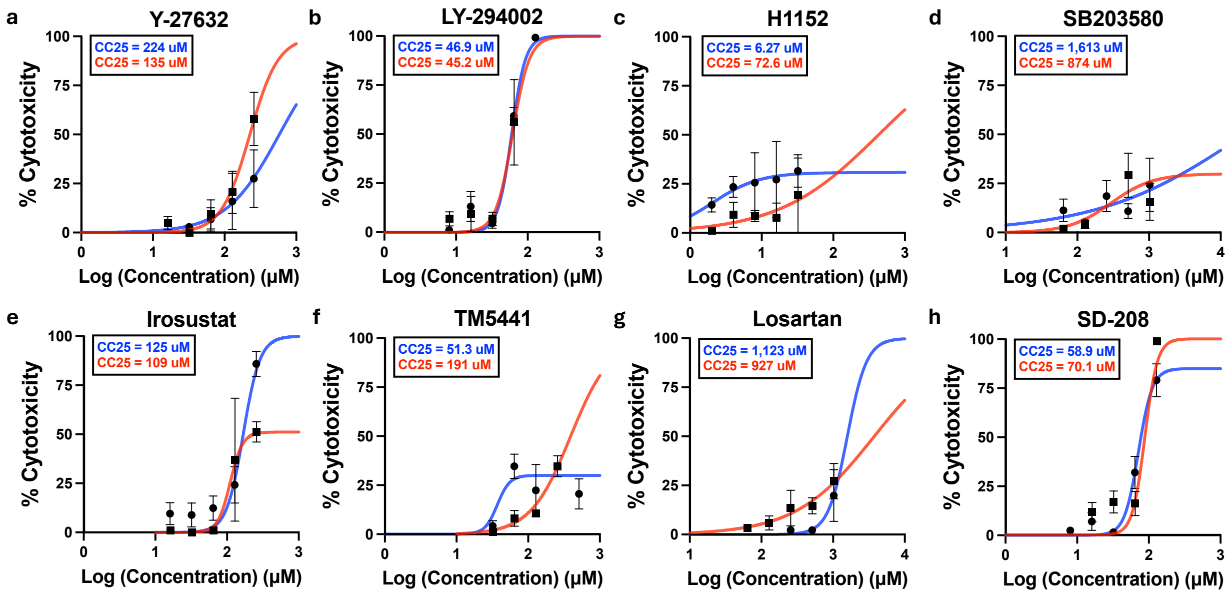


**Supplementary Fig. 7 | Characterizing cytotoxicity of inhibitors in male and female VICs cultured on stiff hydrogels. a-h,** Percent cytotoxicity in male VICs (blue) and female VICs (red) cultured on stiff gels for **a** Y-27632, **b** LY294002, **c** H1152, **d** SB203580, **e** Irosustat, **f** TM5441, **g** Losartan, and **h** SD-208 (n = 3 gels). Data is plotted as mean ± standard error of the mean. The best fit line shown was generated using a nonlinear regression curve fit with GraphPad Prism.

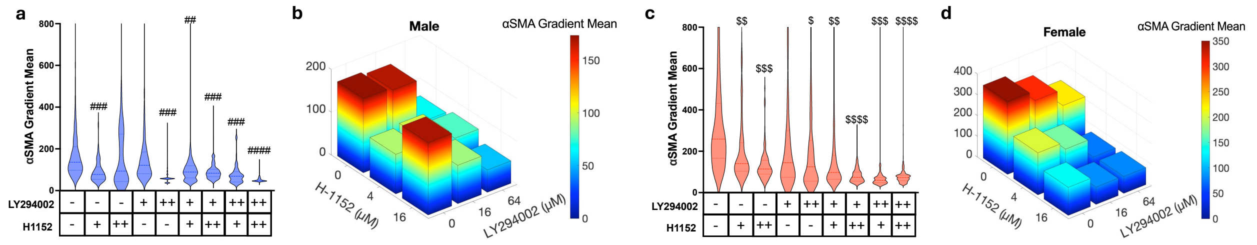


**Supplementary Fig. 8 | The 96-well hydrogel platform captures the combinatorial effects of H-1152 and LY294002 on male and female VICs. a,c,** Violin plot showing the distribution of **a** male or **c** female VIC single cell αSMA gradient mean values for combinations of a low or high dose of H-1152 and/or LY294002 on stiff hydrogels. A minimum of 300 cells were used per condition. **b,d,** 3-D bar plot showing the mean αSMA gradient value for **b** male VICs or **d** female VICs at each experimental condition. - = no drug, + = moderate drug dose, ++ = high drug dose. Statistical significance was determined by one-way ANOVA with Tukey posttests (P<0.0001) and effect size using the Cohen's d-value indicated as ##=d>0.5, ###=d>0.8, ####=d>1.2 relative to male control and $=d>0.2, $$=d>0.5, $$$=d>0.8, $$$$=d>1.2 relative to female control.

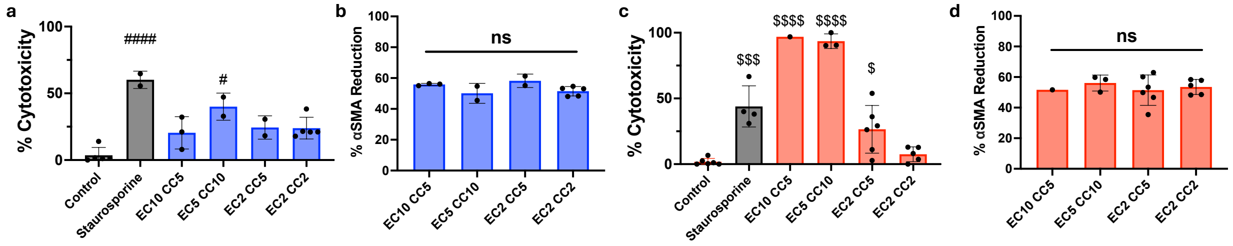


**Supplementary Fig. 9 | A combination of all eight inhibitors at $EC_2/CC_2/10\%\ C_{max}$ doses are non-cytotoxic while still maintaining maximum therapeutic efficacy in male and female VICs cultured on stiff hydrogels. a,c,** Bar graph showing percent cytotoxicity of **a** male or **c** female VICs with different doses of eight drug combinations on stiff hydrogels. **b,d,** Bar graph showing the percent αSMA reduction for **b** male VICs or **d** female VICs with different doses of eight drug combinations on stiff hydrogels (n = 1-6 gels). All drug combinations were additionally limited by 10% $C_{max}$. Data is plotted as mean ± standard deviation. Statistical significance was determined by one-way ANOVA with Tukey posttests. For male VICs, #=P<0.05 and ####=P<0.0001 relative to control. For female VICs, $=P<0.05, $$$=P<0.001, and $$$$=P<0.0001 relative to control.

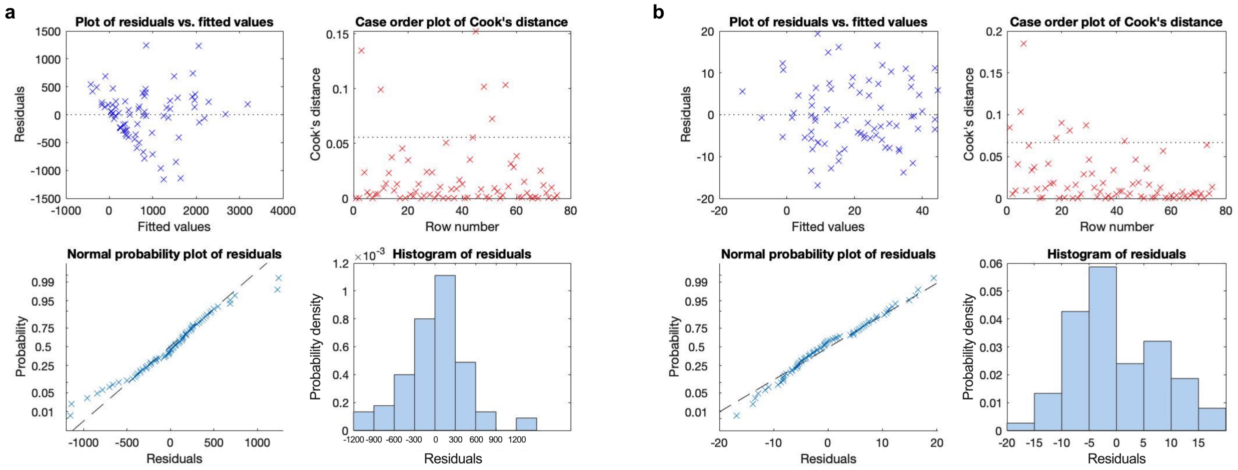


**Supplementary Fig. 10 | Residual-based outlier analysis for percent αSMA reduction data. a,b,** A series of outlier analysis for prospectively acquired percent αSMA reduction data resulting from the 59 OACD combinations for both **a** male and **b** female VICs (n = 2-4 gels). No outliers were detected, and all data were included in the IDentif.AI analysis.

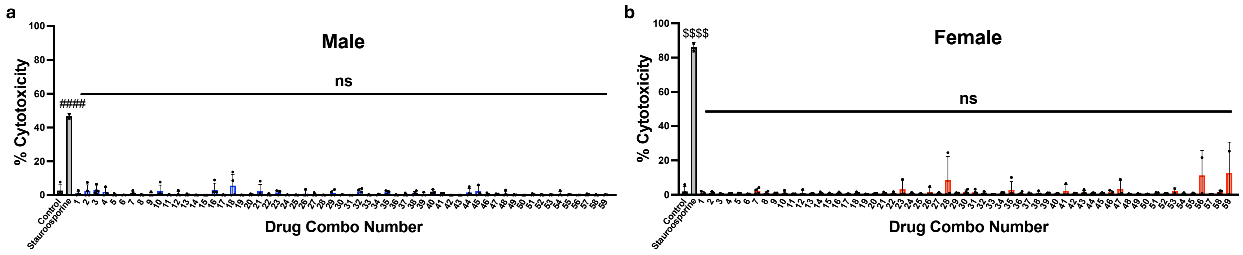


**Supplementary Fig. 11 | Drug combinations used for IDentif.AI analysis of male and female VICs on stiff hydrogels are non-cytotoxic. a,b,** Percent cytotoxicity of all 59 combinations used for IDentif.AI analysis of **a** male VICs or **b** female VICs on stiff hydrogels with positive and negative controls (n = 2-4 gels). Data is plotted as mean ± standard deviation. Statistical significance was determined by one-way ANOVA with Tukey posttests and indicated as ###=P<0.001 relative to male control and $$$$=P<0.0001 relative to female control.

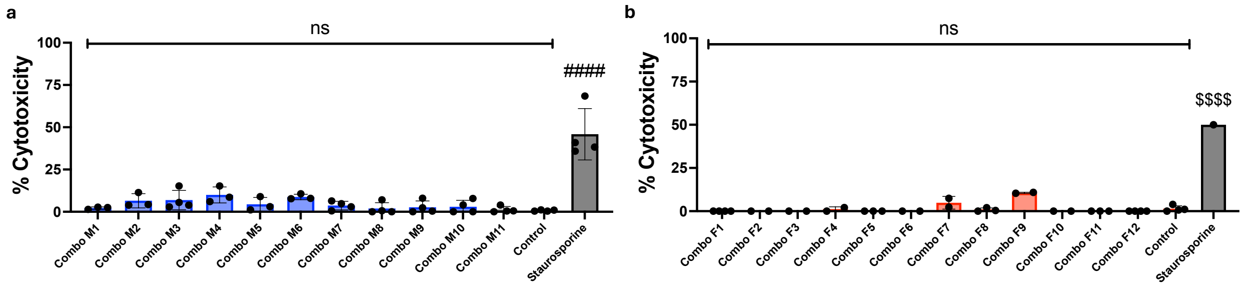


**Supplementary Fig. 12 | Drug combinations used for male and female IDentif.AI validations on stiff hydrogels are non-cytotoxic. a,b,** Percent cytotoxicity of all combinations tested to validate the IDentif.AI model for **a** male VICs or **b** female VICs on stiff hydrogels with positive and negative controls (n = 2-4 gels). Statistical significance was determined by one-way ANOVA with Tukey posttests and indicated as ####=P<0.001 relative to male control and $$$$=P<0.0001 relative to female control.

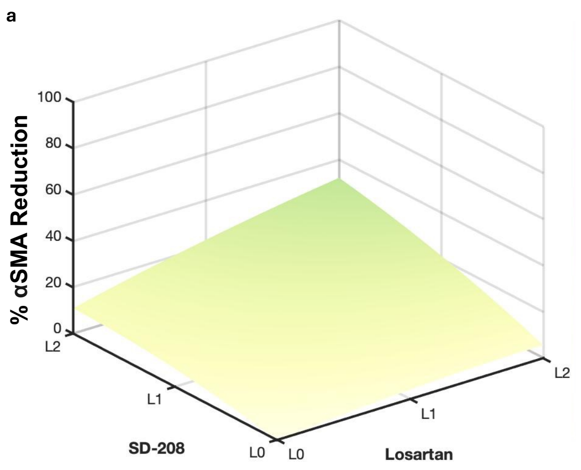


**Supplementary Fig. 13 | Interaction surfaces of IDentif.AI-detected interactions for male and female VIC analyses. a,b,** Interaction surfaces for **a** Losartan/SD-208 and **b** LY294002/H1152 combinations. The predicted interaction of Losartan/SD-208 in male VICs indicated that when both drugs achieve L2 concentrations, the combination may exhibit interactions to enhance efficacy. Similarly, for female VICs, LY294002/H1152 may interact to improve efficacy at L2 concentrations.

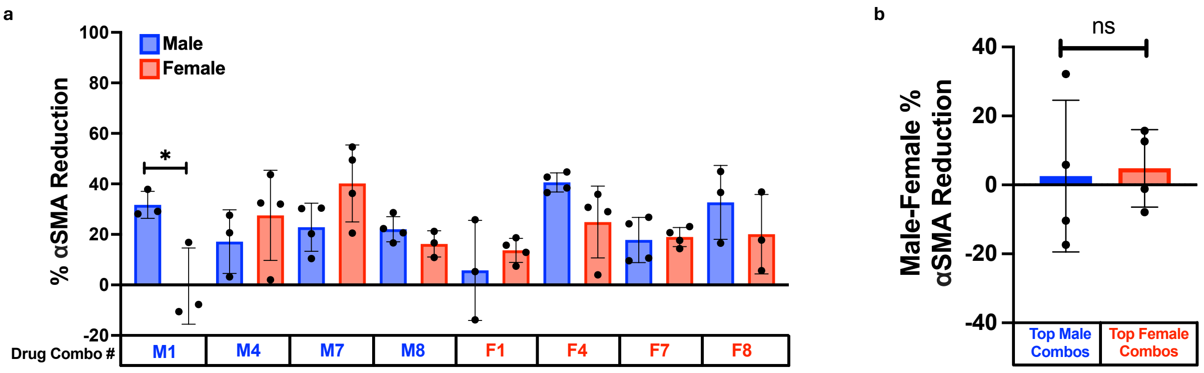


**Supplementary Fig. 14 | Top male and female drug combinations are not sex-specific on stiff hydrogels. a,** Percent αSMA reduction in male and female VICs for top male drug combinations (M1, M4, M7, and M8) and top female drug combinations on stiff hydrogels (F1, F4, F7, and F8) (n = 3-4 gels). **b,** Percent αSMA reduction difference between male and female VICs cultured with top male and female combinations with four drug combinations per group. Data is plotted as mean ± standard deviation. Statistical significance was determined for **a** by one-way ANOVA with Tukey posttests and indicated as *=P<0.05 and for **b** by unpaired two-tailed t-test with Welch's correction.

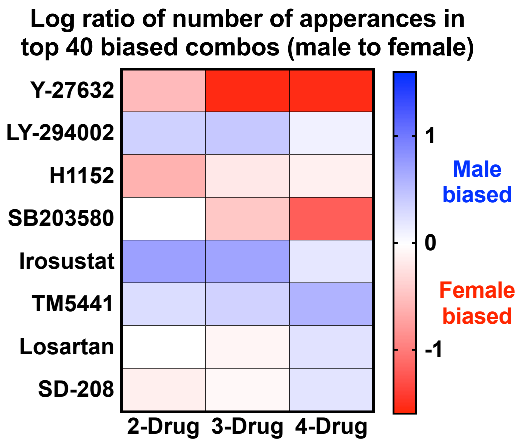


**Supplementary Fig. 15 | Compositions of the top 40 male-biased and female-biased combinations.** Heat map showing the log ratio of the number of drug appearances in the top 40 male-biased combinations to the number of drug appearances in the top 40 female-biased combinations for 2-drug, 3-drug, and 4-drug combinations.

### EXAMPLE CODE

%Load OACD Data and Respective % αSMA reduction Data

data =
[

| Y-27632 | LY-294002 | H-1152 | SB203580 | Irosustat | TM5441 | Losartan | SD-208 | Average |
|---|---|---|---|---|---|---|---|---|
| -1 | -1 | -1 | -1 | -1 | 1 | -1 | -1 | 28.6 |
| 1 | -1 | -1 | -1 | -1 | -1 | 1 | 1 | 40.2 |
| -1 | 1 | -1 | -1 | -1 | -1 | 1 | -1 | 19.2 |
| 1 | 1 | -1 | -1 | -1 | 1 | -1 | 1 | 45.7 |
| -1 | -1 | 1 | -1 | -1 | -1 | -1 | 1 | 27.1 |
| 1 | -1 | 1 | -1 | -1 | 1 | 1 | -1 | 43.3 |
| -1 | 1 | 1 | -1 | -1 | 1 | 1 | 1 | 16.8 |
| 1 | 1 | 1 | -1 | -1 | -1 | -1 | -1 | 25.7 |
| -1 | -1 | -1 | 1 | -1 | -1 | -1 | -1 | 11.1 |
| 1 | -1 | -1 | 1 | -1 | 1 | 1 | 1 | 34.5 |
| -1 | 1 | -1 | 1 | -1 | 1 | 1 | -1 | 11.5 |
| 1 | 1 | -1 | 1 | -1 | -1 | -1 | 1 | 37.2 |
| -1 | -1 | 1 | 1 | -1 | 1 | -1 | 1 | 7.0 |
| 1 | -1 | 1 | 1 | -1 | -1 | 1 | -1 | 3.1 |
| -1 | 1 | 1 | 1 | -1 | -1 | 1 | 1 | 20.0 |
| 1 | 1 | 1 | 1 | -1 | 1 | -1 | -1 | 50.4 |
| -1 | -1 | -1 | -1 | 1 | 1 | 1 | 1 | -7.5 |
| 1 | -1 | -1 | -1 | 1 | -1 | -1 | -1 | 31.5 |
| -1 | 1 | -1 | -1 | 1 | -1 | -1 | 1 | 7.7 |
| 1 | 1 | -1 | -1 | 1 | 1 | 1 | -1 | 0.2 |
| -1 | -1 | 1 | -1 | 1 | -1 | 1 | -1 | 16.2 |
| 1 | -1 | 1 | -1 | 1 | 1 | -1 | 1 | 17.4 |
| -1 | 1 | 1 | -1 | 1 | 1 | -1 | -1 | 54.9 |
| 1 | 1 | 1 | -1 | 1 | -1 | 1 | 1 | 36.5 |
| -1 | -1 | -1 | 1 | 1 | -1 | 1 | 1 | 26.9 |
| 1 | -1 | -1 | 1 | 1 | 1 | -1 | -1 | 42.7 |
| -1 | 1 | -1 | 1 | 1 | 1 | -1 | 1 | 22.4 |
| 1 | 1 | -1 | 1 | 1 | -1 | 1 | -1 | 43.4 |
| -1 | -1 | 1 | 1 | 1 | 1 | 1 | -1 | 41.0 |
| 1 | -1 | 1 | 1 | 1 | -1 | -1 | 1 | 28.4 |
| -1 | 1 | 1 | 1 | 1 | -1 | -1 | -1 | 22.3 |
| 1 | 1 | 1 | 1 | 1 | 1 | 1 | 1 | 43.9 |
| -1 | -1 | -1 | -1 | -1 | -1 | -1 | -1 | -0.2 |
| -1 | 0 | 0 | -1 | -1 | 0 | 1 | 1 | 15.1 |
| -1 | 1 | 1 | -1 | -1 | 1 | 0 | 0 | 24.8 |
| -1 | -1 | 0 | 1 | 0 | -1 | 0 | 0 | 0.7 |
| -1 | 0 | 1 | 1 | 0 | 0 | -1 | -1 | 21.4 |
| -1 | 1 | -1 | 1 | 0 | 1 | 1 | 1 | 13.1 |
| -1 | -1 | 1 | 0 | 1 | -1 | 1 | 1 | 4.3 |
| -1 | 0 | -1 | 0 | 1 | 0 | 0 | 0 | -0.7 |
| -1 | 1 | 0 | 0 | 1 | 1 | -1 | -1 | 19.0 |
| 0 | -1 | 0 | 0 | -1 | 0 | -1 | 0 | 26.1 |
| 0 | 0 | 1 | 0 | -1 | 1 | 1 | -1 | 20.7 |
| 0 | 1 | -1 | 0 | -1 | -1 | 0 | 1 | 35.4 |
| 0 | -1 | 1 | -1 | 0 | 0 | 0 | 1 | 18.0 |
| 0 | 0 | -1 | -1 | 0 | 1 | -1 | 0 | 32.5 |
| 0 | 1 | 0 | -1 | 0 | -1 | 1 | -1 | 40.1 |
| 0 | -1 | -1 | 1 | 1 | 0 | 1 | -1 | 21.4 |
| 0 | 0 | 0 | 1 | 1 | 1 | 0 | 1 | 29.8 |
| 0 | 1 | 1 | 1 | 1 | -1 | -1 | 0 | 40.8 |
| 1 | -1 | 1 | 1 | -1 | 1 | -1 | 1 | 6.4 |
| 1 | 0 | -1 | 1 | -1 | -1 | 1 | 0 | 24.7 |
| 1 | 1 | 0 | 1 | -1 | 0 | 0 | -1 | 22.1 |
| 1 | -1 | -1 | 0 | 0 | 1 | 0 | -1 | 25.0 |
| 1 | 0 | 0 | 0 | 0 | -1 | -1 | 1 | 17.3 |
| 1 | 1 | 1 | 0 | 0 | 0 | 1 | 0 | 46.0 |
| 1 | -1 | 0 | -1 | 1 | 1 | 1 | 0 | -6.2 |
| 1 | 0 | 1 | -1 | 1 | -1 | 0 | -1 | 28.1 |
| 1 | 1 | -1 | -1 | 1 | 0 | -1 | 1 | 29.3 |

]

%Define Inputs and Outputs

```
x = data(: , 1:8);
y = data(: , 9);

%IDentif.AI Analysis

result = stepwiselm(x , y, ‘quadratic’, 'ResponseVar', 'Inhibition', 'PredictorVars', {‘Y-27632’, ‘LY-294002’, ‘H-1152’, ‘SB203580’, ‘Irosustat’, ‘TM5441’, ‘Losartan’, ‘SD-208’});
```
